# A non-canonical Hippo pathway regulates spindle disassembly and cytokinesis during meiosis in *Saccharomyces cerevisiae*

**DOI:** 10.1101/2020.02.21.959619

**Authors:** Scott M. Paulissen, Cindy A. Hunt, Christian J. Slubowski, Yao Yu, Dang Truong, Xheni Mucelli, Hung T. Nguyen, Shayla Newman-Toledo, Aaron M. Neiman, Linda S. Huang

## Abstract

Meiosis in the budding yeast *Saccharomyces cerevisiae* is used to create haploid yeast spores from a diploid mother cell. During meiosis II, cytokinesis occurs by closure of the prospore membrane, a membrane that initiates at the spindle pole body and grows to surround each of the haploid meiotic products. Timely prospore membrane closure requires *SPS1*, which encodes a STE20-family GCKIII kinase. To identify genes that may activate *SPS1*, we utilized a histone phosphorylation defect of *sps1* mutants to screen for genes with a similar phenotype and found that *cdc15* shared this phenotype. *CDC15* encodes a Hippo-like kinase that is part of the mitotic exit network. We find that Sps1 complexes with Cdc15, that Sps1 phosphorylation requires Cdc15, and that *CDC15* is also required for timely prospore membrane closure. We also find that *SPS1*, like *CDC15*, is required for meiosis II spindle disassembly and sustained anaphase II release of Cdc14 in meiosis. However, the NDR-kinase complex encoded by *DBF2/DBF20 MOB1* which functions downstream of *CDC15* in mitotic cells, does not appear to play a role in spindle disassembly, timely prospore membrane closure, or sustained anaphase II Cdc14 release. Taken together, our results suggest that the mitotic exit network is rewired for exit from meiosis II, such that *SPS1* replaces the NDR-kinase complex downstream of *CDC15*.

## INTRODUCTION

Sexual reproduction requires meiosis for the production of haploid gametes from a diploid precursor cell. The events of meiosis such as spindle disassembly and cytokinesis must be properly coordinated with each other, and with the developmental events that occur during gametogenesis. A better understanding of how these events are coordinated is important for understanding gamete formation.

In the budding yeast *Saccharomyces cerevisiae*, the haploid gametes are spores, which form when diploid cells encounter starvation conditions where nitrogen and carbon are limiting (reviewed in Neiman 2011). During sporulation, the diploid mother cell remodels its interior to form four haploid spores. Spore morphogenesis begins with the formation of a prospore membrane that grows from the spindle pole body. The prospore membranes grow around the haploid nuclei and fuse to close at the side of the nucleus away from the spindle pole body, resulting in the capture of each nucleus within its own membrane (Diamond *et al*. 2009). A protein complex known as the Leading Edge Protein complex is at the growing edge of the prospore membrane and includes Ssp1, Ady3, Irc10, and Don1 (Knop and Strasser 2000; Moreno-Borchart *et al*. 2001; Nickas and Neiman 2002; Maier *et al*. 2007; Lam *et al*. 2014).

Prospore membrane closure is the cytokinetic event in meiosis, and involves the removal of the Leading Edge Protein complex (Maier *et al*. 2007). Proper timing of prospore membrane closure requires *SPS1,* which encodes a STE20-family GCKIII kinase; cells lacking *SPS1* produce hyperelongated prospore membranes that close later than those in wild-type cells (Slubowski *et al*. 2014; Paulissen *et al*. 2016). Prospore membrane closure must be properly coordinated with other meiosis II events, such as spindle disassembly.

Compared to meiosis, exit from mitosis, which involves the downregulation of CDK activity and the coordination of spindle disassembly and cytokinesis, has been more extensively studied. Mitotic exit involves the activation of the Tem1-GTPase at the spindle pole body as it moves into the newly formed bud, leading to the activation of the Cdc15 Hippo-like kinase (Mah *et al*. 2001; Visintin and Amon 2001; D’Aquino *et al*. 2005; Pereira and Schiebel 2005; Maekawa *et al*. 2007; Chan and Amon 2010; Rock and Amon 2011; Bertazzi *et al*. 2011; Falk *et al*. 2016). Cdc15 phosphorylates the spindle pole body localized Nud1 scaffold, which leads to the recruitment and activation of the NDR kinase complex, Dbf2-Mob1 (Gruneberg *et al*. 2000; Luca *et al*. 2001; Rock and Amon 2013). A decrease in mitotic cyclin dependent kinase (CDK) activity is also required for Cdc15 and Mob1 activation (Campbell *et al*. 2019). Activation of the NDR kinase complex promotes the sustained release of the Cdc14 serine-threonine phosphatase from the nucleolus to inactivate mitotic CDK activity and promote exit from mitosis (Visintin *et al*. 1998; Shou *et al*. 1999; Mohl *et al*. 2009; Manzoni *et al*. 2010). These components are part of the Mitotic Exit Network (MEN) (reviewed in Bardin and Amon 2001; Stegmeier and Amon 2004; Hergovich and Hemmings 2012; Weiss, 2012; Juanes and Piatti 2016).

Meiotic exit has been shown to utilize some but not all of the MEN components. Exit from meiosis I does not require the MEN (Kamieniecki *et al*. 2005; Pablo-Hernando *et al*. 2007; Attner and Amon, 2012), which instead acts to coordinate exit from meiosis II. *CDC15* plays a role in meiosis II spindle disassembly (Pablo-Hernando *et al*. 2007; Attner and Amon 2012) and is also required to maintain nuclear and nucleolar release of Cdc14 in meiosis II (Pablo-Hernando *et al*. 2007). Furthermore, a prospore membrane closure (Diamond *et al*. 2009) and morphology (Pablo-Hernando *et al*. 2007) defect have been described for *cdc15*. However, the upstream MEN component *TEM1* does not appear to play a role in Cdc15 activation, as the Tem1-GTPase is not seen at the spindle pole body in meiosis (Attner and Amon 2012) and Tem1-depleted cells complete meiosis with similar efficiencies as wild-type cells (Kamieniecki *et al*. 2005). The spindle pole body located scaffold encoded by *NUD1* is also likely not involved in exit from meiosis, as *nud1* temperature sensitive alleles do not disrupt meiosis (Gordon *et al*. 2006) and *NUD1* is not required for Dbf20 kinase activity in meiosis (Attner and Amon 2012).

In meiosis, the NDR-kinase complex utilizes the Mob1 regulatory subunit along with either of the paralogous Dbf20 and Dbf2 NDR kinases (Attner and Amon 2012; Renicke *et al*. 2017). *MOB1* plays a role in meiosis II, as *mob1* cells progress through meiosis I with wild type kinetics, but show a delay in exit from meiosis II (Attner and Amon, 2012). Dbf20 kinase is active in meiosis II, and its kinase activity as well as its interaction with the Mob1 regulatory subunit is dependent on *CDC15* in meiosis II, although deletion of *DBF20* did not show a delay in meiosis II exit (Attner and Amon 2012). The major phenotype seen for cells lacking the NDR kinases complex in meiosis is a defect in spore number control (Renicke *et al*. 2017); spore number control is a process that involves the selection of nuclei associated with younger spindle pole bodies over older spindle pole bodies for spore packaging when available energy resources are a low (Davidow *et al*. 1980; Nickas *et al*. 2004; Taxis *et al*. 2005). Nud1 is also involved in spore number control (Gordon *et al*. 2006; Renicke *et al*., 2017).

Here, we examine timely prospore membrane closure, meiosis II spindle disassembly and Cdc14 sustained release in anaphase II and find that *CDC15* and *SPS1* act together to regulate exit from meiosis II. However, the NDR kinase complex encoded by *DBF2 DBF20 MOB1* does not seem to be involved in these events. Instead, *DBF2 DBF20 MOB1* are important for spore number control, as previously demonstrated (Renicke *et al*. 2017). Likewise, we find that *CDC15* and *SPS1* are not involved in controlling spore number and appear to act separately from the NDR kinase complex in meiosis II.

## MATERIALS AND METHODS

### Yeast strains, growth and sporulation

All strains used in this study are in the SK1 background (Kane and Roth 1974) and are described in Supplemental Material Tables S1 and S2. All strains are derived from LH177 (Huang *et al*. 2005) except for YS429 (see below), the previously published strains (AN117-4B, A20239, A22416, and HI50), and the published strains used for screening (see below and Supplemental Table 1 and 3); alleles from these strains were crossed into the LH177 derived SK1 strain background. Standard genetic methods were used to create and propagate strains unless otherwise noted (Rose and Fink 1990). Epitope-tagged strains and knock out alleles were created using PCR-mediated recombination methods, as previously described (Longtine *et al*. 1998; Lee *et al*. 2013; Slubowski *et al*. 2015).

YS429 was constructed by replacing the native *DBF2* promoter with the *CLB2* promoter by PCR mediated integration using pRK67 (Kaminiecki *et al*. 2005) as a template in strain AN117-4B (Neiman *et al*. 2000). The resulting haploid was crossed to a *dbf20Δ::kanMX6* haploid from the yeast knockout collection (Rabitsch *et al*. 2001) and segregants from this cross were mated to create YS429. The *CDC15-9MYC* allele in LH1070 and LH1071 was from A22416 (Attner *et al*. 2012). The *mob1-mn* (*KanMX6:pCLB2-3HA-MOB1*) allele used in this study is from A20239 (Attner *et al*. 2012). The *cdc15-mn* (*mxKAN:prCLB2:HA:CDC15*) allele used in this study is from HI50 (Pablo-Hernando *et al*. 2007).

Unless otherwise noted, cells were grown in standard yeast media and sporulated in a synchronous manner in liquid media, as previously described (Huang *et al*. 2005). In brief, liquid cultures were grown with agitation at 30°C. Cells to be sporulated were first grown to saturation in YPD overnight at 30° and then transferred to YPA and grown to ∼1.5 OD600/ml overnight. These cells were then harvested, washed in double-distilled H2O (ddH2O), and resuspended in 1% potassium acetate (KOAc) at an OD600/ml of 2.0. Sporulation of cells containing plasmids was the same as above except instead of YPD, cells were grown in synthetic dextrose (SD) media, lacking the appropriate nutrient for selection.

### Plasmids

The plasmid pRS426-E20 was created by PCR amplification of GFPEnvy from pFA6a-link-GFPEnvy-SpHIS5 (Slubowski *et al*. 2015) using primers OLH1669 (GTGTggatccATGTCTAAAGGCGAGGAATTG) and OLH1679 (GTGTgaattcTTTGTACAATTCGTCCATTCCTAA), which incorporated the *Bam*HI and *Eco*RI restriction sites flanking GFPEnvy. The amplified fragment was then digested with *EcoRI* and *Bam*HI. pRS426-G20 (Nakanishi *et al*. 2004) was also digested with *EcoRI* and *Bam*HI, removing the GFP from in front of the *SPO20* fragment on that plasmid. The resulting linearized backbone was then ligated to the GFPEnvy PCR fragment. The resulting plasmid was verified by sequencing.

### Screening for H4S1p phenotype

To screen for a H4S1p phenotype, mutant strains were inoculated in 20 ml YPD and grown overnight. Cultures were then diluted 1:100 into 80 ml YPA, such that the OD600 was between 0.1-0.2, and grown overnight to reach an OD600 between 1.0-1.2. Cells were then collected, washed in ddH2O, and resuspended in 50 ml of 2% KOAc at an OD600 of 1.2 (∼2×107 cells/ml). 10 mls of cells were collected at 0, 8, 10, and 24 hours after induction of sporulation. Proteins were extracted by resuspending cells in Breaking Buffer (50 mM Tris-HCL pH7.5, 10% glycerol, 1 mM EDTA, 10mM MgCl2, 100mM NaCl, 1 mM DTT) with protease inhibitors (1 mM PMSF, 1 µg/ml leupeptin, and 1 µg/ml peptastin A) and phosphatase inhibitors (100 mM NaF, 100 mM Na4P2O7, 10 mM Na3VO4). Cells were lysed using glass beads and a bead-beater. Protein concentration of extracts was determined using the Bio Rad Protein Assay, and extracts were adjusted to similar concentrations. Loading buffer was added to extract, which were then boiled and loaded onto an SDS-PAGE gel and immunoblotted. H4S1p was detected using a rabbit anti-phospho H4/H2A S1p antibody (07-179; Upstate/Merck-Millipore) at a dilution of 1:4000, detected using HRP-conjugated secondary antibodies and ECL reagents (Amersham/GE Healthcare), and exposed to X-ray film.

### Immunoblotting

For all immunoblotting experiments other than those performed for the H4S1p screening, cells were collected at the indicated times and prepared using the TCA precipitation method (Philips and Herskowitz 1998), which involves first lysing cells in a lysis buffer (1.85 N NaOH and 10% v/v β-mercaptoethanol) followed by precipitation of proteins with 50% (v/v) trichloroacetic acid (TCA). Precipitated protein lysates were then washed with ice-cold acetone and resuspended in 2× sample buffer neutralized with 5 µl of 1 M Tris base; samples were heated before loading. Protein lysates were separated on standard single percentage SDS-PAGE gels, except for the histone phosphorylation blot in Figure 1A, which was run on a Novex 10-20% Tricine gel (Invitrogen).

**Figure 1.**
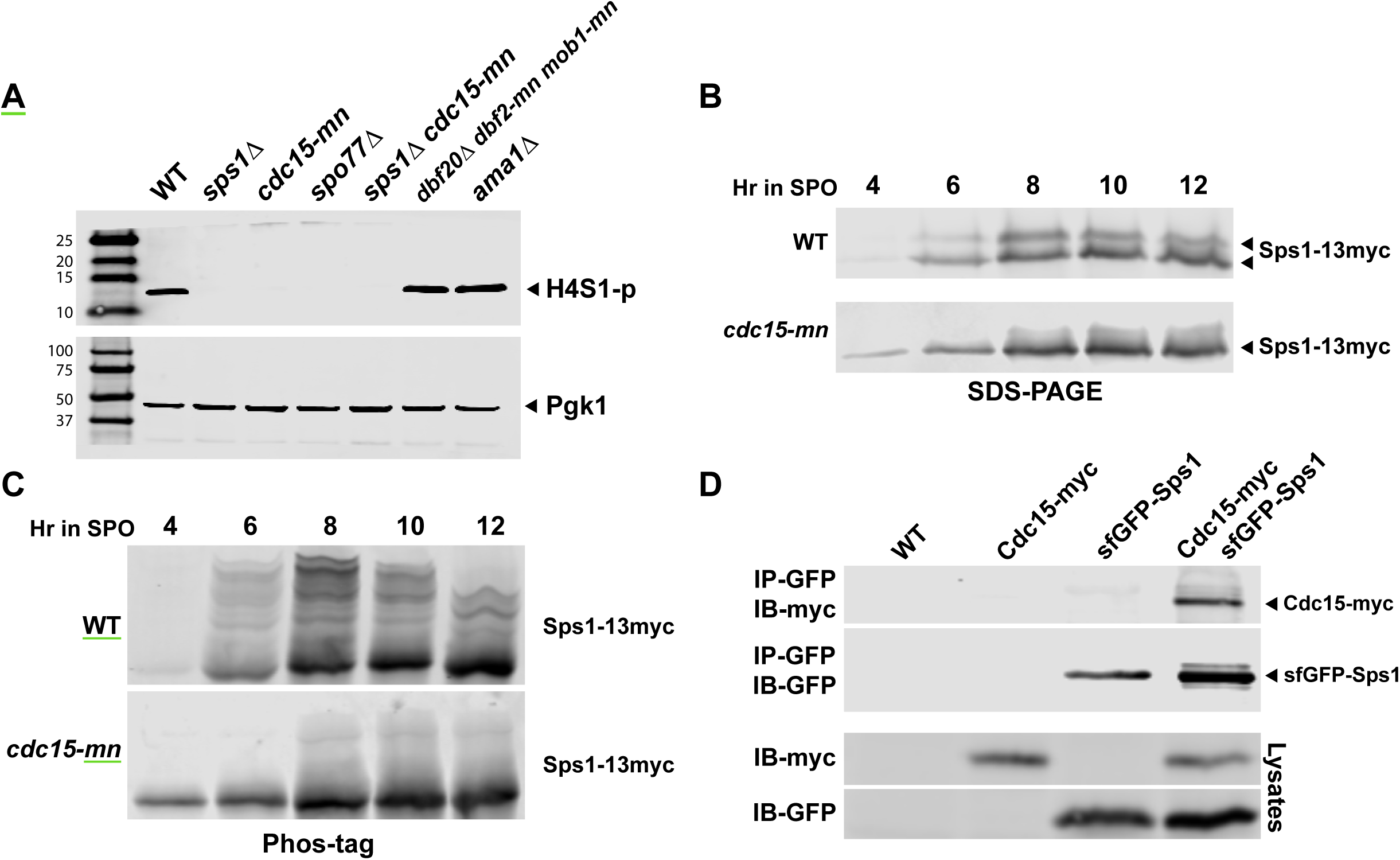
*CDC15* is required for *SPS1* phosphorylation. (A) Screening for other genes deficient in Histone phosphorylation. Cells lacking specific genes were induced to sporulated and collected at 8 hours after induction of sporulation. H4S1 phosphorylation was assayed by immunoblotting. Pgk1 was used as a loading control and was from the top half of the same gel as that probed for histone phosphorylation. Protein marker sizes shown to the left of gel. Wild type (WT (LH902)), *sps1* (LH966), *cdc15* (LH1066), *spo77* (LH1010)*, sps1 cdc15* (LH1067), *mob1 dbf2 dbf20* (LH1068), *ama1* (LH1014). (B) Sps1-13myc was assayed on an SDS-PAGE gel using lysates from WT (LH875)) and *cdc15-mn* (LH1069) cells that were collected at the indicated times after induction of sporulation and probed with an anti-myc antibody. (C) Sps1-13myc was assayed using a Phos-tag gel using lysates from the same samples collected for (B). (D) Cdc15 and Sps1 form a complex. Immunoprecipitation experiments were carried out using lysates from WT (LH902), *Cdc15-myc* (LH1070), *sfGFP-Sps1* (LH986), *Cdc15-myc sfGFP-Sps1* (LH1071). Sps1 was immunoprecipitated (IP) using GFP-Trap beads. Immunoblots (IB) were probed with either anti-GFP antibody or anti-myc antibody.

The separated protein extract was transferred onto Immobilon LF-PVDF membrane, blocked with LI-COR PBS block, and incubated with the appropriate primary antibodies. H4S1 phosphorylation was detected using the anti-phospho histone H4/H2A S1p antibody at 1:1000 (Upstate/Merck-Millipore); sf-GFP-Sps1 was detected using JL-8 anti-GFP antibodies (Takara/Clontech) at 1:1000; Sps1-13x*myc* and Cdc15-9x*myc* were detected using 9E10 anti-myc antibodies (Covance) at 1:1000; Pgk1 was detected by using 22C5D8 anti-Pgk1 (Life Technologies) at 1:1000; Fluorescent infrared-dye-conjugated anti-mouse secondary antibodies were used at 1:10,000 (LI-COR). All membranes were imaged using an Odyssey Infrared Imaging System (LI-COR).

### Immunoprecipitation

Lysates for immunoprecipitation were prepared from 120 OD600 of cells. Cell pellets were lysed in a MiniBeadBeater8 (Biospec) at 4°C with glass beads in IP buffer (300mM NaCl, 5 mM EGTA (pH 8.0), 50 mM Tris (pH 7.4), and 0.5% Nonidet P-40) with added protease and phosphatase inhibitors as previously described (Huang *et al*. 2005).

Lysate was clarified with three spins at maximum speed in a tabletop microcentrifuge, and an aliquot was saved for examination by immunoblot; this aliquot was first TCA precipitated before loading onto an SDS-PAGE gel. For immunoprecipitation, clarified lysate was then added to 40μl of blocked agarose beads (ChromoTek) incubated on a nutator at 4°C for 30 minutes. Lysates incubated on a nutator at 4°C for two hours with 20μl of GFP-Trap beads (ChromoTek). GFP-Trap complexes were then washed four times in IP buffer and re-suspended in 2× SDS-PAGE sample buffer, boiled for 5 minutes, clarified through centrifugation and then separated by SDS-PAGE.

### Phos-tag analysis

Phos-tag gels were made using Phos-tag acrylamide (WACO) at a final concentration of 31.4 μM Phos-tag and 50.6 μM MnCl2 in an otherwise standard 6% SDS-polyacrylamide gel, as described in (Whinston *et al*. 2013). Samples were prepared as above and run at 80 V at 4°C before being transferred and imaged, as above.

### Microscopy

Widefield Microscopy was performed using a 100x (NA 1.45) objective on a Zeiss Axioskop Mot2. Images were taken using an Orca-ER cooled CCD camera (Hamamatsu) using Openlab 4.04 (Perkin Elmer) or iVision (BioVision Technologies) for image acquisition. Confocal Microscopy was performed using a 100x (NA 1.49) objective on a Zeiss LSM-880 Confocal Microscope. Confocal images were acquired using Zeiss Zen-Black software. Images were cropped and merged using ImageJ and FIJI (Schneider *et al*. 2012; Schindelin *et al*., 2012).

### Assaying prospore membrane closure, formation, and number

Cells were assayed for prospore membrane closure and formation as previously described (Paulissen *et al*. 2016). For prospore membrane closure and formation, only cells in anaphase II (as determined by Htb2-mCherry) were counted. Cells were considered to have initiated prospore membranes if a single prospore membrane could be detected. Cells were considered to have closed their prospore membranes if a single rounded prospore membrane was detected within the ascus.

To assay the number of prospore membranes that form within the mother cell, cells were sporulated in 1% acetate and fixed using 4.5% methanol free formaldehyde. Only cells in anaphase II (as determined by Htb2-mCherry) were counted. Cells were counted on a Zeiss Axioskop Mot2 using a 100x (NA 1.45) objective. Strains were sporulated in triplicate; 100 anaphase II cells were counted per culture, for a total of 300 cells per strain.

### Visualization of spindles by immunostaining

Sporulating cells were harvested and fixed in 3.7% methanol-free formaldehyde for 15 minutes, washed twice with ddH_2_O and then suspended in 1 ml of SP buffer (1M sorbitol 10mM pH7.8 PBS). These cells were then spheroplasted at 37°C after adding 20μl 20T zymolyase at 5 mg/ml concentration and 1 μl β-mercaptoethanol; spheroplast formation was checked using a 100x phase objective on a Zeiss AxioMot2. Spheroplasted cells were washed with 1 ml SP buffer and resuspended in 500 μl of fresh SP buffer. Spheroplasted cells were then adhered to polylysine coated slides, blocked with blocking buffer (1% BSA, 0.1% Triton in PBS) and then rinsed three times with PBS.

Tubulin was detected using monoclonal mouse 12G10 anti-Tub1 antibody at 1:1000 concentration (Developmental Studies Hybridoma Bank). Cells were incubated with antibody for one hour, rinsed four times with PBS and then incubated with Cy2-conjugated donkey anti-mouse antibodies at 1:100 dilution for one hour (JacksonImmuno; Figure 2) or AlexaFluor 488 conjugated Donkey anti-mouse at 1.25 mg/ml (JacksonImmuno; Figure 8). Stained cells were then washed three times with PBS, twice with ddH_2_O and then sealed in Vectashield mounting medium (Vector Labs).

**Figure 2.**
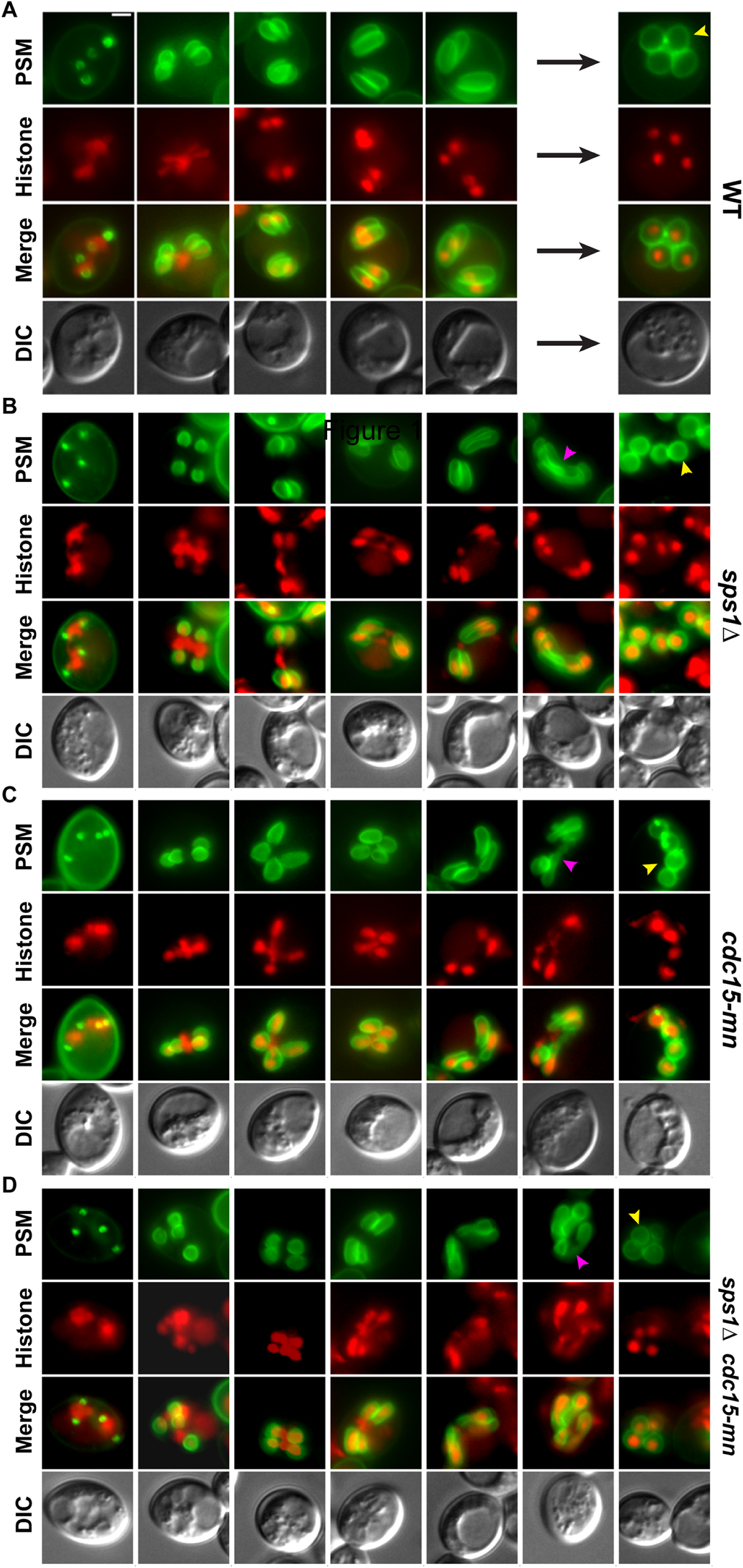
*CDC15* is required for proper prospore membrane development. Prospore membranes are labelled in green using the plasmid pRS426-G20 (WT (LH917) and *sps1Δ* (LH1047) or pRS426-E20 (*cdc15-mn* (LH1073), *sps1Δ cdc15-mn* (LH1074). Histones are labeled in red using genomically integrated *HTB2-mCherry* fusion protein. Developmental stages are shown from early (left) to late (right). Pink arrowheads point to examples of hyperelongated prospore membranes. Yellow arrowheads point to examples of rounded prospore membranes. Images were captured using a wide-field microscope.

### Statistical analysis

Statistical comparisons were performed by 1-way ANOVA followed by Tukey HSD post hoc test. All tests were performed using JMP11 (SAS).

### Data availability

The strains and plasmids created for this study are available upon request. Supplemental Figures and Tables are available at FigShare. The data necessary for confirming the conclusions of this article are present.

## RESULTS

### *CDC15* is required for Sps1 phosphorylation

Phosphorylation of the Ser1 residue of Histone H4 is greatly increased during meiosis and Sps1 had previously been demonstrated to be important for this phosphorylation (Krishnamoorthy et al., 2006). To identify additional genes that may function with Sps1, we used a Western blot assay with a H4/H2A Serine1 phosphorylation (H4/H2A S1p)-specific antibody to initially screen through a few genes involved in sporulation (*ama1*, *cdc15*, *gip1*, *spo71*, *spo73*, *spo75*, *spo77*, and *ssp2*) for those that display an H4 phosphorylation defect similar to *sps1Δ* mutants. We then carried out a more unbiased screen, examining H4 phosphorylation in a subset of strains from a collection of mutants in genes that are upregulated in sporulation (Rabitsch *et al*. 2001). The 120 genes that were tested are listed in Supplemental Material Table S3.

*cdc15* and *spo77* were among the mutants identified in this screen that exhibited decreased histone phosphorylation similar to *sps1Δ* (Figure 1A). *SPO77* was isolated as a high copy suppressor of a hypomorphic allele of *sps1* and acts with *SPS1* in regulating timely prospore membrane closure (Paulissen *et al*. 2016). Because a link between *SPS1* and *CDC15* was not previously reported, we focused our studies on *CDC15*.

Since Sps1 is a phosphoprotein (Slubowski *et al*. 2014) and because *CDC15* encodes a Hippo-like protein kinase (Schweitzer and Philippsen, 1991; Rock *et al*. 2013), we asked whether *CDC15* was required for Sps1 phosphorylation. We examined Sps1 phosphorylation state in sporulating cells with depleted levels of Cdc15. Separation of Sps1 on an SDS-PAGE gel revealed that the doublet seen in wild type cells (Slubowski *et al*. 2014) collapses into the faster migrating band in the *cdc15-mn* strain (*cdc15-m*eiotic *n*ull; *CDC15* under the control of the mitotic *CLB2* promoter (Lee and Amon 2003; Pablo-Hernando *et al*. 2007)) (Figure 1B). This result suggested that post-translational modification of Sps1 protein was *CDC15* dependent.

To better examine the migration shifts due to post-translational phosphorylation, we used a Phos-tag polyacrylamide gel to resolve the Sps1 protein. Phos-tag gels specifically retard the migration of phosphorylated protein species through the gel (Kinoshita *et al*. 2006). Sps1 runs as multiple bands on a Phos-tag gel, consistent with it being a phosphoprotein (Figure 1C). This banding pattern was strikingly reduced in the *cdc15-mn* strain (Figure 1C), which supports the idea that *CDC15* is required for most, if not all, of the phosphorylation of Sps1.

To determine whether the phosphorylation of Sps1 by Cdc15 may be direct, we examined whether Cdc15 and Sps1 physically interact in sporulating cells by co-immunoprecipitation. Using protein lysates from a strain containing both *CDC15-13myc* and *sfGFP-SPS1*, we see Cdc15 and Sps1 in a complex (Figure 1D).

Because Cdc15 is a phosphoprotein (Jaspersen and Morgan 2000; Jones *et al*. 2011), we asked if post-translational modifications of Cdc15 were altered in *sps1Δ* mutants. SDS-PAGE analysis of Cdc15 in both wild-type and *sps1Δ* mutant cells both show a distinct doublet suggesting phosphorylation of Cdc15 is not altered in the *sps1*Δ mutant (Figure S1), consistent with *CDC15* acting upstream of *SPS1*. Taken together, these results show that *CDC15* is required for Sps1 phosphorylation and support a model in which Cdc15 is the upstream activating kinase of Sps1.

### Like *SPS1*, *CDC15* is required for timely prospore membrane closure

Previous studies have demonstrated a role for *CDC15* in prospore membrane morphogenesis, with *cdc15* mutant cells forming aberrant prospore membrane morphologies (Pablo-Hernando *et al*. 2007) and having a defect in closing prospore membranes (Diamond, et al. 2009). To visualize prospore membranes, we utilized GFP (either eGFP or GFPEnvy, a bright and photostable GFP variant (Slubowski *et al*. 2015)) fused to the 40 amino acid prospore membrane-targeting region of the Spo20 protein (Nakanishi *et al*. 2004). We examined prospore membranes in live cells during sporulation in wild type, *sps1*Δ cells and *cdc15-mn* cells. Unlike wild type cells (Figure 2A), *cdc15-mn* cells show hyperelongated prospore membranes (Figure 2B), similar to those see in *sps1*Δ cells (Figure 2C), consistent with the previously described *cdc15* prospore membrane morphology (Pablo-Hernando *et al*. 2007) and closure defect (Diamond *et al*. 2009).

We asked whether *SPS1* and *CDC15* acted in the same or in a parallel pathway, to regulate prospore membrane closure. We created the *sps1Δ cdc15-mn* strain, and saw that the double mutant cells displayed a prospore membrane morphology defect that was no worse than that of the *sps1*Δ mutation alone (Figure 2C and 2D), consistent with both genes acting in the same pathway.

Because *SPS1* plays a role in timely prospore membrane closure (Paulissen *et al*. 2016), we asked whether *CDC15* affects the timing of prospore membrane closure. To assay prospore membrane closure, we examined the appearance of rounded prospore membranes, as rounded prospore membranes appear when the membrane closes (Diamond *et al*. 2009; Paulissen *et al*. 2016). Cells with *cdc15-mn* exhibited both a delay in appearance of as well as a reduction in the accumulation of closed prospore membranes, forming rounded membranes at approximately 72% (Figure 2C; Figures 3A), similar to the reduction seen in *sps1Δ* mutants and less than the 95% seen in wild type cells (Figure 3A and Paulissen *et al*. 2016). The observed delay in prospore membrane closure is not due to a delay in prospore membrane initiation, as *cdc15-mn* cells showed a similar onset of prospore membrane biogenesis as wild type (Figure 3B), similar to the lack of defect seen in prospore membrane initiation seen in *sps1Δ* mutants (Figure 3B, Paulissen *et al*. 2016).

**Figure 3.**
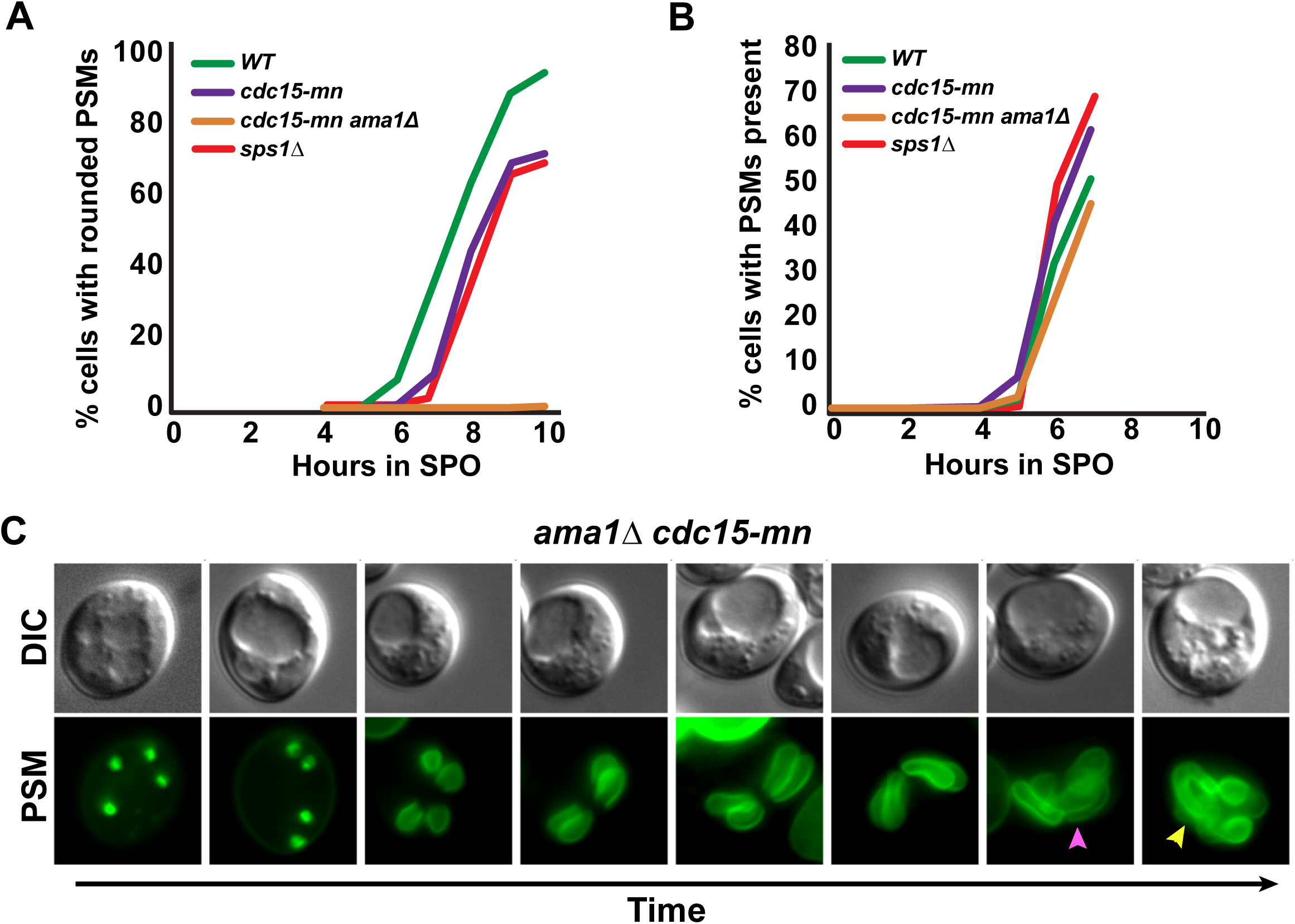
*CDC15* and *SPS1* are required for timely prospore membrane closure and act in parallel to *AMA1*. Quantitation of prospore membrane (PSM) closure (A) and initiation (B) in WT (LH917), *cdc15-mn* (LH1072), *sps1Δ* (LH1047) *cdc15-mn ama1Δ* (LH1075). At least 100 cells were counted per timepoint, for each genotype. Prospore membranes were visualized using the plasmid pRS426-G20 for the WT and *sps1Δ* strains and pRS426-E20 for the *cdc15-mn* and *ama1Δ cdc15-mn* strains. (C) *cdc15-mn ama1*(LH1076) mutants produce hyperelongated prospore membranes that do not close. Prospore membranes are labelled in green using the plasmid pRS426-E20. Prospore membranes are shown from early (left) to late (right) on the bottom row, with a corresponding DIC picture of the cell on top. Pink arrowheads point to examples of hyperelongated prospore membranes. Yellow arrowheads point to examples of rounded prospore membranes. Images were captured using a wide-field microscope.

*SPS1* acts to regulate timely prospore membrane closure in a pathway in parallel to *AMA1*, as cells lacking *SPS1* or *AMA1* have partial defects in prospore membrane closure that is exacerbated in the double mutant (Paulissen, et al. 2016). We tested whether *CDC15* also acts in parallel to *AMA1* and examined doubly mutant *cdc15-mn ama1Δ* cells. We found that *cdc15-mn ama1Δ* cells form rounded prospore membranes at < 0.5% frequency (Figure 3A), a much stronger defect than either *cdc15-mn* (Figure 3A) or *ama1Δ* cells alone (∼30%; Diamond et al. 2009; Paulissen et al. 2016). These *cdc15-mn ama1Δ* double mutant cells form prospore membranes that become highly invaginated, filling the cytoplasmic space of the mother cell and only rarely rounding up and closing (Figure 3C), similar to that seen in the *sps1Δ ama1Δ* double mutant (Paulissen *et al*. 2016). These results taken together show that *CDC15* regulates timely prospore membrane closure, acting in the same pathway as *SPS1* and in parallel to *AMA1*.

### *SPS1* has a meiosis II spindle disassembly defect similar to *CDC15*

Cells lacking *CDC15* have been previously shown to have a meiosis II spindle disassembly defect (Pablo-Hernando, *et al*. 2007; Attner and Amon 2012). Since *SPS1* and *CDC15* share prospore membrane phenotypes, we examined whether *SPS1* played a role in meiotic spindle disassembly.

We examined spindles in wild type, *sps1Δ* and *cdc15-mn* sporulating cells by immunostaining fixed sporulating cells. Spindles in wild type cells elongated and then disassembled during meiosis I and II, eventually forming small spindles in the newly created spores (Figure 4A). *cdc15-mn* cells failed to disassemble meiosis II spindles, with late anaphase II spindles becoming extended and ultimately fragmenting within the cell (Figure 4B), consistent with previous observations (Pablo-Hernando *et al*. 2007; Attner and Amon 2012).

**Figure 4.**
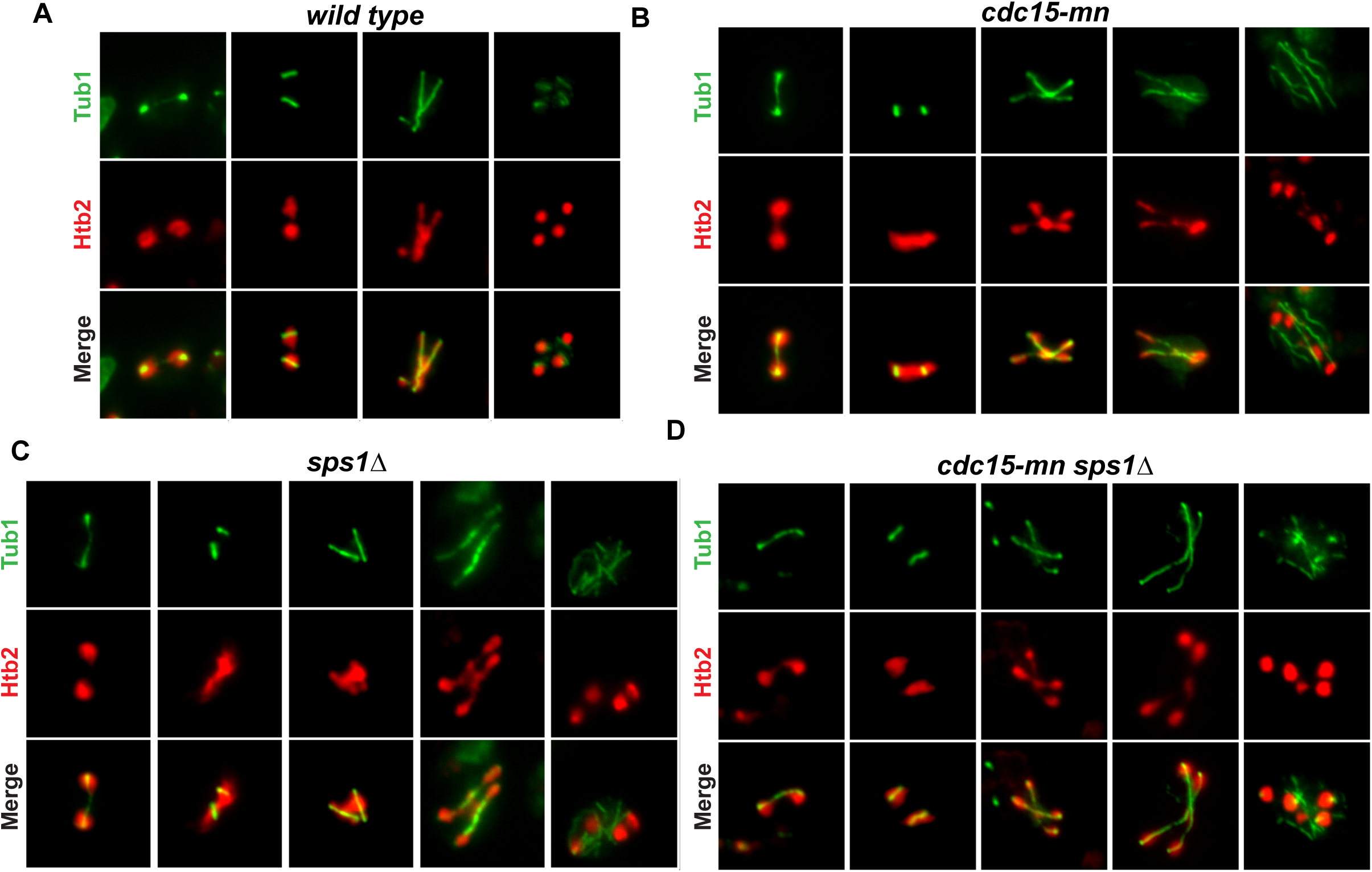
*SPS1* has a spindle disassembly defect. Microtubules were visualized in green using an anti-Tub1 antibody. Histones, in red, are visualized using *HTB2-mCherry*. Cells at different time points in meiosis, arrayed from early (left) to late (right), with varying terminal phenotypes shown for the mutant strains. Cells were fixed at appropriate times during sporulation and stained with anti-Tub1 antibodies. Images were captured using a wide-field microscope. Cells are of the following genotypes: (A) WT (LH902) (B) *cdc15-mn* (LH1072) (C) *sps1Δ* (LH976) (D) *cdc15-mn sps1Δ* (LH1067).

*sps1Δ* mutant cells had microtubule morphologies that were indistinguishable from that of the *cdc15-mn* mutant (Figure 4C), including the frequent occurrence of elongated, fragmented, and supernumerary microtubules late in anaphase II (Figure 4C). When we examine the meiotic spindles in the *sps1Δ cdc15-mn* double mutant, we see that the microtubule phenotype was indistinguishable to that of the single mutants (Figure 4D). These results are consistent with *SPS1* and *CDC15* acting in the same pathway for meiotic exit, which involves both meiotic spindle disassembly and cytokinesis, the latter accomplished via prospore membrane closure during yeast meiosis.

### Cdc14 sustained release in anaphase II requires *SPS1*

During mitosis, the MEN, a signal transduction network that utilizes Cdc15 activation of Dbf2-Mob1 NDR kinase complex (Rock et al. 2013), will ultimately promote the release of the Cdc14 phosphatase from the nucleolus to inactivate mitotic CDK activity and promote exit from mitosis (Visintin *et al*. 1998; Shou *et al*. 1999; Mohl *et al*. 2009; Manzoni *et al*. 2010). In meiosis, MEN is thought to be predominately active in meiosis II, with Dbf20 as the major NDR kinase in meiosis, although Dbf2 also plays a role (Attner and Amon 2012; Renicke et al. 2017).

During meiosis, *CDC14* acts in both meiosis I and meiosis II (Buonomo et al. 2003; Marston et al. 2003; Kamieniecki et al. 2005; Villoria et al. 2016; Fox et al., 2017). In meiosis, Cdc14 is released from the nucleolus before anaphase I spindle elongation, then reappears in the nucleolus at the start of meiosis II and is released again just before anaphase II (Bizzari and Marston 2011; Kerr et al. 2011); the initial release of Cdc14 in meiosis requires the FEAR network and not the MEN (Buonomo *et al*. 2003; Marston *et al*. 2003; Kamieniecki *et al*. 2005; Pablo-Hernando *et al*. 2007). However, *CDC15* is required for the sustained release of Cdc14 during anaphase II (Pablo-Hernando et al. 2007; Attner and Amon, 2012).

We first re-examined Cdc14 release during anaphase II in wild type cells, using a *CDC14-GFPEnvy* allele. We see dynamic localization for Cdc14 (Figure 5), as previously described (Bizzari and Marston 2011), with Cdc14 being released from the nucleolus and into the nucleus and cytoplasm during anaphase II. We also see, as previously described (Pablo-Hernando *et al*. 2007), that Cdc14 release is not properly sustained in anaphase II in the *cdc15-mn* mutants.

**Figure 5.**
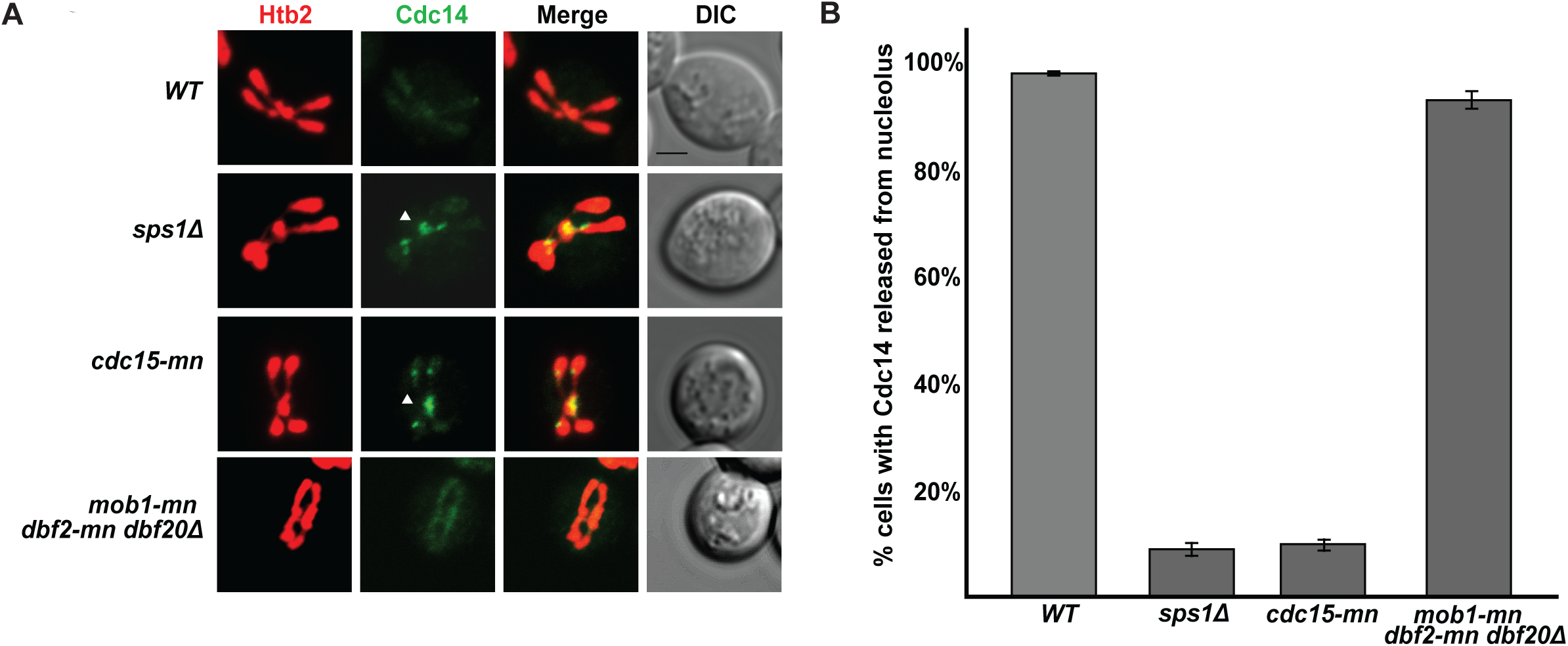
The sustained release of Cdc14 requires *SPS1* and *CDC15* but not *DBF2 DBF20 MOB1*. (A) The Cdc14-GFPEnvy fusion protein was visualized in WT (LH1077), *sps1Δ* (LH1078), *cdc15-mn* (LH1079) and *mob1-mn dbf2-mn dbf20Δ* (LH1080) cells. Representative images are shown from these strains. Histones are visualized using a genomically integrated *Htb2-mCherry*. Images were captured using a confocal microscope. White arrowhead points to nucleolar-localized Cdc14. Scale bar = 2 µm. (B) Quantitation of cells in anaphase II (as determined by Htb2-mCherry localization) with Cdc14 released from the nucleolus. Cells were sporulated in triplicate, with 100 anaphase II cells counted for each biological replicate for a total of 300 cells per strain. Error bars represent standard error of the mean. The wild type and triple mutant (*mob1-mn dbf2-mn dbf20Δ*) strains are significantly different from the *cdc15-mn* and the *sps1Δ* strains, but not from one another (one-way ANOVA [F(3,8)=860, p<0.001], followed by Tukey HSD post hoc test (alpha = 0.01)).

Given the role of *CDC15*, we asked whether *SPS1* plays a role in Cdc14 anaphase II release. When we examined Cdc14 release from the nucleolus in *sps1Δ* cells during anaphase II, we see that Cdc14 release is not properly sustained, similar to that seen in *cdc15-mn* mutants (Figure 5). We confirmed localization of the Cdc14 to the nucleolus in *sps1Δ* and *cdc15-mn* mutants using the nucleolar marker Nop56/Sik1 (Gautier *et al*. 1997; Figure S2).

Because the Dbf2-Mob1 NDR kinase complex acts in between *CDC15* and *CDC14* during mitosis, we examined the role of NDR kinase complex in Cdc14 release in anaphase II. We created the *dbf2-mn* allele, which places the mitotically-required *DBF2* gene under the control of the mitosis-specific *CLB2* promoter. To eliminate as much NDR kinase complex activity in meiosis as possible, we combined the *dbf2-mn* allele with the previously constructed *mob1-mn* and the *dbf20Δ* alleles (Attner and Amon 2012). When we examined Cdc14 release in the *mob1-mn dbf2-mn dbf20Δ* triple mutant strain, we find that Cdc14 is properly released during anaphase II, similar to what is seen in wild type cells and in contrast to what is seen in the *cdc15* and *sps1* mutant cells (Figure 5). Thus, in meiosis II, the NDR kinase complex, encoded by *MOB1 DBF2 DBF20*, does not act downstream of *CDC15* to regulate Cdc14 release. Instead, our results are consistent with *SPS1* acting downstream of *CDC15* to regulate Cdc14 sustained release during anaphase II.

### *CDC15* and *SPS1* do not act with the NDR kinase complex for spore number control

The NDR kinase complex has been previously shown to play a role in spore number control, a process that determines the number of spores packaged during meiosis (Renicke *et al*. 2017). Spore number control regulates the number of spindle pole bodies that are competent for prospore membrane growth; this process depends on a spindle pole body modification that happens based on the age of the spindle pole body and the nutrients available to sporulating cells (Davidow *et al*. 1980; Nickas *et al*. 2004; Taxis *et al*. 2005). Depletion of the NDR kinase complex results in fewer spores per ascus forming during sporulation, as seen when *MOB1 DBF2 DBF20* activity was reduced using a protein depletion system (Renicke *et al*. 2017). We see a similar result using our *mob1-mn dbf2-mn dbf20Δ* strain, as assayed by counting refractile spores formed (Figure S3) or by counting the number of prospore membranes formed as a proxy for the number of spores than can form within the ascus (Figure 6).

**Figure 6.**
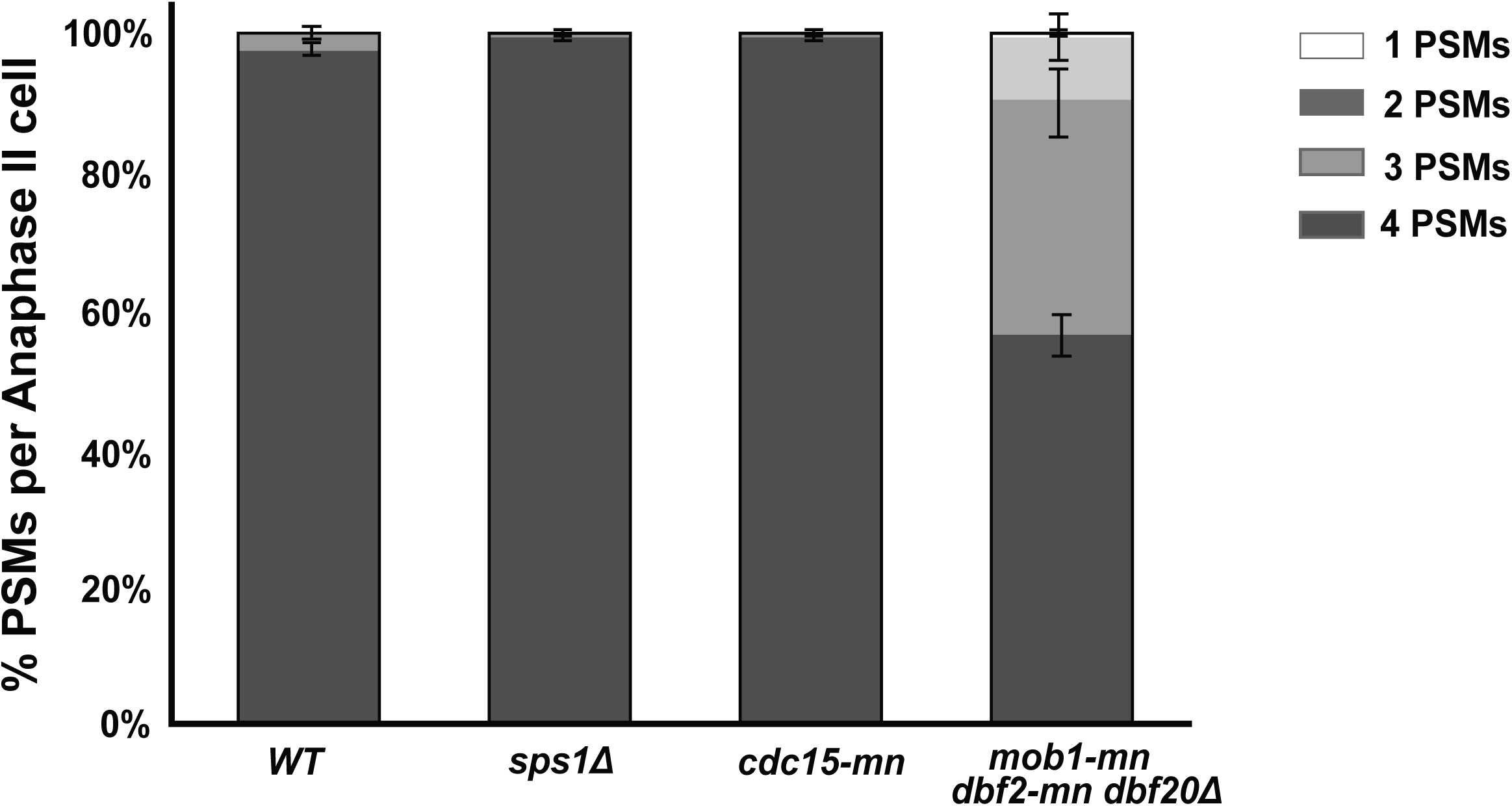
*SPS1* and *CDC15* are not required to regulate the number of prospore membranes formed. The number of prospore membranes formed per cell were counted in anaphase II cells, as assayed by visualizing histones using *Htb2-mCherry*. Prospore membranes were visualized using the plasmid pRS426-E20. WT (LH1081), *sps1Δ* (LH1089), *cdc15-mn* (LH1073), *mob1-mn dbf2-mn dbf20Δ* (LH1082). Three biological replicates of 100 cells per replicate were counted, for a total of 300 cells per strain. Error bars represent standard error of the mean. The *wild type*, *cdc15-mn* and *sps1Δ* strains are significantly different from the triple mutant (*mob1-mn dbf2-mn dbf20Δ*) strain, but not from one another, using 4 PSMs as the variable for comparison (one-way ANOVA [F(3,8) = 437, p<0.001], followed by Tukey HSD post hoc test (alpha = 0.01)).

Because neither *cdc15-mn* nor *sps1Δ* cells form refractile spores, we assayed spore number control by counting the number of prospore membranes that are present in anaphase II, to determine how many spores could form within an ascus. We see most *sps1Δ* and *cdc15-nm* mutant cells will initiate four prospore membranes per ascus, similar to that seen in wild type cells, and unlike that seen in the *mob1-mn dbf2-mn dbf20Δ* mutants. These results suggest that neither *sps1Δ* nor *cdc15-nm* act with the NDR kinase complex in spore number control.

### The NDR kinase complex does not play a role in timely prospore membrane closure or spindle disassembly

Because we see that the Mob1-Dbf2/20 NDR kinase complex appears to regulate distinct biological processes from the Cdc15 and Sps1 kinases, we examined prospore membrane morphology and timing of prospore membrane closure in the *mob1-mn dbf2-mn dbf20Δ* triple mutant. When we examine prospore membrane morphology in the *mob1-mn dbf2-mn dbf20Δ* triple mutant, we do not see the characteristic hyperelongated prospore membranes seen in *sps1Δ* and *cdc15-nm* mutant cells, although aberrant prospore membrane size and nuclear capture defects were observed (Figure 7A). When we examined the timing of prospore membrane closure, we saw that the *mob1-mn dbf2-mn dbf20Δ* mutant cells produced rounded prospore membranes with similar timing as wild type cells and do not exhibit the delay seen in *cdc15-mn* or *sps1Δ* mutant cells (Figure 7A and Figure B); all cells examined initiate formation of prospore membranes at a similar time (Figure 7C).

**Figure 7.**
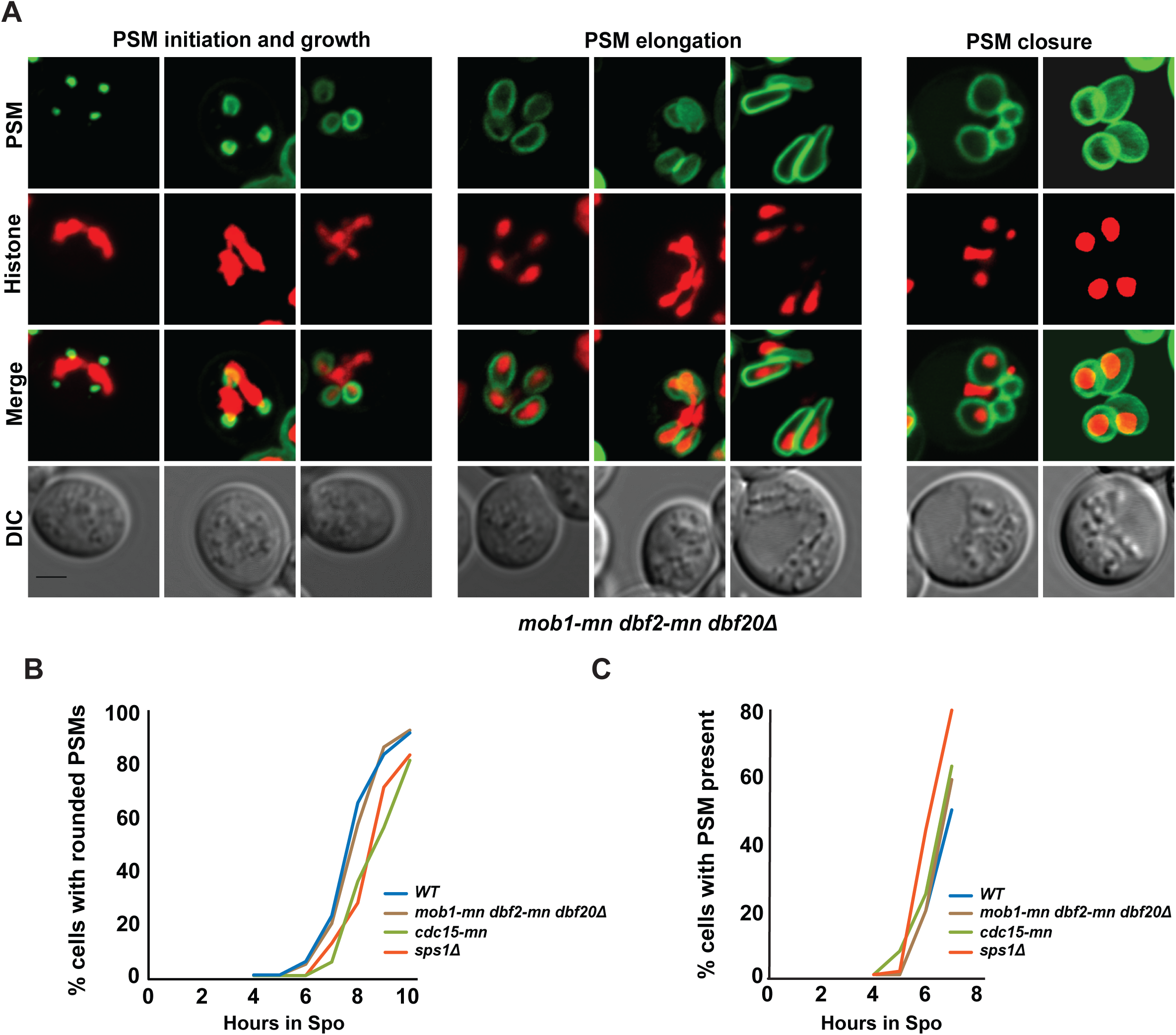
*DBF2 DBF20 MOB1* are not required for timely prospore membrane closure. (A) *mob1-mn dbf2-mn dbf20Δ* do not form hyperelongated prospore membranes (PSMs). Prospore membranes are labelled in green using the plasmid pRS426-E20. Histones are visualized using *HTB2-mCherry*. Scale bar = 2µm. *mob1-mn dbf2-mn dbf20Δ* cells close (B) and initiate (C) prospore membranes with timing similar to wild type cells. Prospore membrane closure and initiation were counted as in Figure 3A and B, with at least 200 cells counted per timepoint for each genotype. WT (LH1081), *mob1-mn dbf2-mn dbf20Δ* (LH1082), *sps1Δ* (LH1089), and *cdc15-mn* (LH1073); the pRS426-E20 plasmid was transformed into these strains for visualization of the prospore membrane.

Because we see a spindle disassembly defect in *sps1* and *cdc15-mn* mutant cells, we examined the spindle in the *mob1-mn dbf2-mn dbf20Δ* cells. *mob1-mn dbf2-mn dbf20Δ* triple mutant cells do not produce the elongated, fragmented, and supernumerary microtubules late in anaphase II that are seen in the *sps1Δ*, *cdc15-nm*, and *sps1Δ cdc15-nm* double mutant cells. Instead, in late meiosis II, spindles in the *mob1-mn dbf2-mn dbf20Δ* cells appear to be disassembled into shorter punctate pieces (Figure 8A), which is distinct from the fragmented microtubules seen in *sps1Δ*, *cdc15-nm*, and *sps1Δ cdc15-nm* late in anaphase II. Thus, the NDR kinase complex does not appear to play a role in timely prospore membrane closure or spindle disassembly.

**Figure 8.**
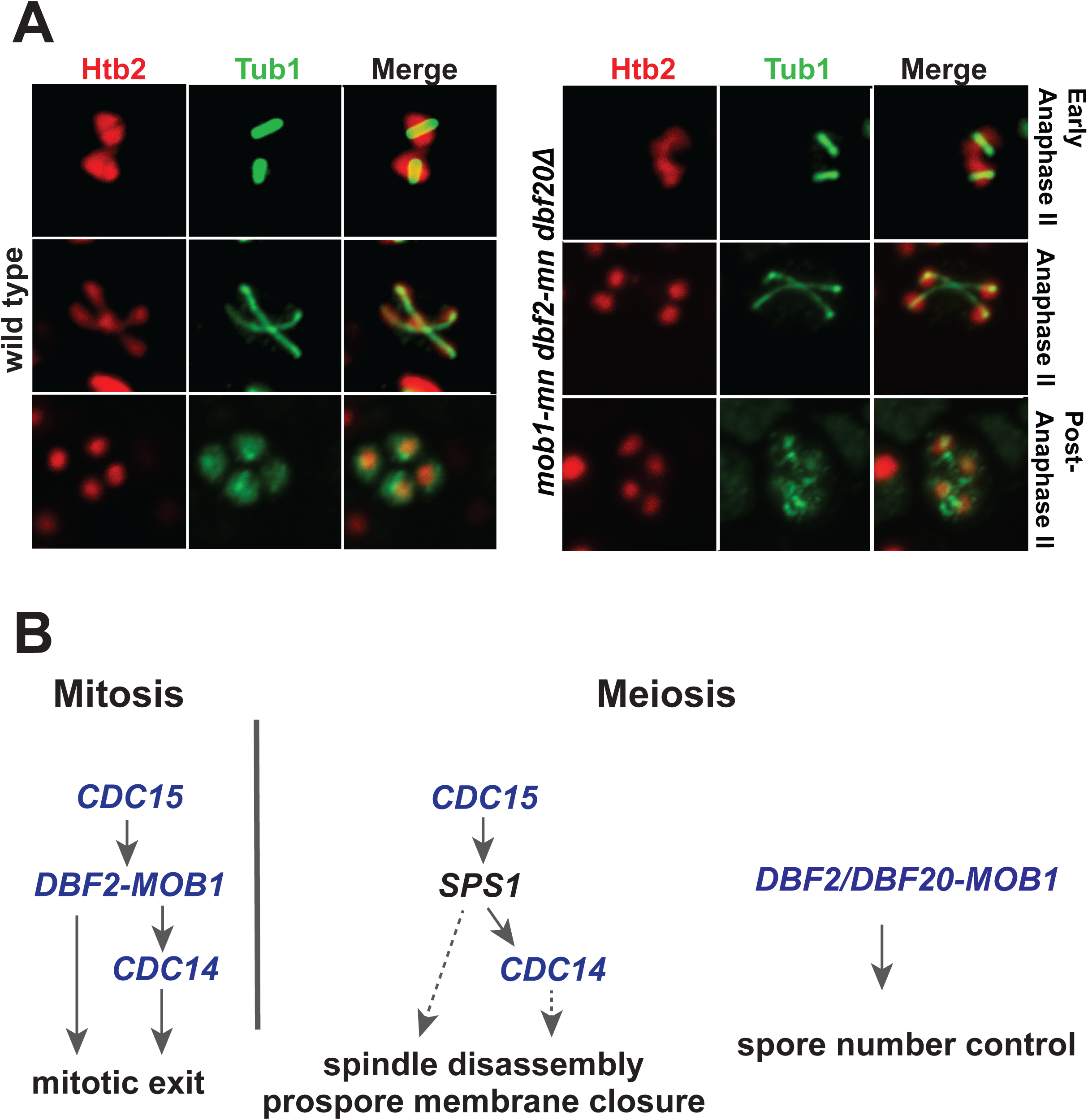
*DBF2 DBF20 MOB1* are not required for spindle disassembly. (A) Spindles, as seen in wild type (LH902) and *mob1-mn dbf2-mn dbf20Δ* (LH1068) cells. Cells were fixed and stained. Microtubules were visualized in green using an anti-Tub1 antibody. Histones, in red, are visualized using *HTB2-mCherry*. Nuceli were visualized using *HTB2-mCherry*. (B) Model depicting the relationship between mitotic exit members in mitosis and meiosis. See discussion in <u>text.</u>

## DISCUSSION

Our studies demonstrate that during meiosis, timely prospore membrane closure, meiosis II spindle disassembly, and sustained release of Cdc14 at anaphase II are regulated by *SPS1* and *CDC15*, while the Mob1-Dbf2/20 complex plays a separate role in meiosis regulating spore number control. These results suggest that for exit from meiosis II, the MEN is rewired, such that Sps1 replaces the NDR kinase complex and acts downstream of the Cdc15 kinase.

### *SPS1* acts with *CDC15* to regulate exit from meiosis II

We describe two previously unknown roles for *SPS1* in the completion of meiosis: timely spindle disassembly and Cdc14 sustained release. Prior to this study, the involvement of *SPS1* in sporulation was thought to be for spore morphogenesis (Friesen *et al*. 1994; Iwamoto *et al*. 2005) and more specifically, for timely prospore membrane closure (Paulissen *et al*. 2016). Furthermore, *sps1Δ* and *cdc15-mn* mutants have identical phenotypes, as we describe a role for *CDC15* in timely prospore membrane closure. Since we see that Cdc15 is needed for Sps1 phosphorylation, these results are consistent with a model where Sps1 acting downstream of Cdc15 for exit from meiosis II (see model in Figure 8B).

A better understanding of the mechanism underlying how this pathway leads to the exit of meiosis II will require identification of downstream targets. In mitosis, although the phosphorylation of many CDK targets are reversed by Cdc14 upon mitotic exit, some downstream targets important for cytokinesis are directly phosphorylated by the Dbf2 kinase (Meitinger *et al*., 2011, Oh *et al*., 2012). For meiosis, it is unknown whether all targets downstream of *CDC15* and *SPS1* are directly regulated by the Cdc14 phosphatase, or whether Sps1 may directly phosphorylate downstream targets as well. It is likely that Sps1 plays some direct role, as previous studies have demonstrated that although *CDC15* is required for sustained Cdc14 release, *CDC14* does not appear to play a role in meiosis II spindle disassembly or prospore membrane morphology (Pablo-Hernando et al. 2007; Arguello-Miranda et al. 2017). These studies depleted *CDC14* activity using a *cdc14-ΔNES* allele which deleted the Cdc14 nuclear export signal at residues 359-367 (Pablo Hernando et al. 2007) or the *cdc14-3* ts allele (Arguello-Miranda et al., 2017). The role of *SPS1* in prospore membrane closure is likely to be *CDC14* independent, as *SPS1* is required for the phosphorylation and reduced stability of Ssp1 (Paulissen *et al*. 2016), a protein localized to the leading edge of the growing prospore membrane that must be removed and degraded for this process to occur (Maier *et al*. 2007).

We find that *CDC15* and *SPS1* act in parallel to *AMA1*, which encodes a meiosis-specific activator of the anaphase promoting complex (APC/C) (Cooper *et al*. 2000). Previous studies have examined a hyperactive *ama1* allele (*ama1-m8*, which lacks eight consensus Cdc28 phosphorylation sites in Ama1) in combination with *cdc15-mn* and found a significant increase in prospore membrane closure in the double mutant (Diamond *et al*. 2009), consistent with our findings here. Interestingly, *AMA1* has also been linked to both spindle disassembly and prospore membrane closure. For meiosis II spindle disassembly, *AMA1* acts downstream of *HRR25* encoded casein kinase 1 (Arguello-Miranda *et al*. 2017). *AMA1* regulates prospore membrane closure (Diamond *et al*. 2009; Paulissen *et al*. 2016) and affects the stability of Ssp1, localized at the leading edge of the prospore membrane (Diamond *et al*. 2009). The combination of both meiosis II spindle disassembly and prospore membrane closure defects for *cdc15, sps1,* and *ama1* mutants raises the question of whether the prospore membrane closure defect seen in these mutants is a consequence of the stable meiosis II spindles, which are in the way and thus prevent the membrane fusion event required to close the membrane. Whether prospore membrane closure and spindle disassembly are coordinated by the regulation of a common target of both these pathways, or, whether these two events are regulated via distinct targets remains to be determined.

### Cdc15 and the NDR/LATS kinase complex play distinct roles in meiosis

In meiosis II, cells appear to utilize *CDC15* and *MOB1-DBF2/20* for distinct roles, unlike in mitosis where Cdc15 activates a conserved Mob1-NDR kinase signaling system, as seen in typical Hippo signaling (Hergovich and Hemmings, 2012, Weiss, 2012). In meiosis II, it appears that *MOB1-DBF2/20* is important for spore number control (Renicke et al., 2017), in which neither *CDC15* nor *SPS1* play a role, as assayed by the number of prospore membranes formed.

Previous work described a role for *CDC15* in spore number control, with *cdc15* depleted mutants forming more meiotic plaques on the spindle pole bodies when sporulated in low acetate conditions, compared to wild type cells and the *mob1 dbf2 dbf20* triple mutant (Renicke *et al*. 2017). We do not see a difference between *cdc15-mn* and wild type cells in spore number control using a direct assay of counting the number of prospore membranes formed in 1% acetate (Figure 6). Under our sporulation conditions, it may not be possible to see the *CDC15* effect, as most wild type cells produce four prospore membranes (although we can see the effect of the NDR/LATS kinase complex on spore number control under these conditions (Figure S3)). Importantly, the previous study found that the *mob1 dbf2 dbf20* depleted triple mutant had a distinct phenotype from *cdc15* depleted mutants in spore number control (Reinicke *et al*. 2017), consistent with our findings that Cdc15 and the NDR/LATS kinase complex play distinct roles in meiosis (Figure 8B).

Previous studies have shown that Dbf20 kinase activity depends on *CDC15* in meiosis II, and the interaction of Dbf20 and Mob1 is dependent on *CDC15* (Attner and Amon 2012). However, our phenotypic characterization is consistent with the exit from meiosis functions of *CDC15* not requiring *DBF2/20-MOB1*. It is possible that there are some Dbf20-Mob1 functions in meiosis that require *CDC15* activity, as the kinase activity of Dbf20 and its interaction with Mob1 were demonstrated biochemically.

### GCK kinase as an alternative member of the Hippo signaling pathway

Sps1 is a STE20-family GCKIII kinase (Slubowski *et al*. 2014), and modifications of the typical Hippo signaling module to include STE20-family GCK kinases has been reported. For example, in fission yeast, Hippo signaling also involves in intervening GCK-family kinase, Sid1, that acts between the Cdc7 Hippo-like kinase and the Mob1/Sid2 NDR kinase for septation (referred to as the SIN pathway) (Guertin et al., 2000). For tracheal morphogenesis in *Drosophila*, the NDR kinase Trc is activated by Germinal center kinase III, a GCKIII kinase, (Poon et al., 2018). Unlike these previously described cases of GCK use that involve a downstream NDR/LATS kinase, for budding yeast meiosis, it appears that there has been a separation of function between the Hippo-GCKIII module and the downstream Mob1-Dbf2/20 NDR/LATS kinase, providing a distinct example of how Hippo signaling can act with GCK members.

## Acknowledgements

We thank the Amon lab for strains, Yasuyuki Suda for assistance with strain construction, Matt Durant for comments on the manuscript, and Angelika Amon for comments on an earlier version of this work. This work was funded by grants from NIH to LSH (GM86805) and AMN (GM072540) and a Sanofi-Genzyme Fellowship to SMP.

## REFERENCES

Argüello-Miranda, O., I. Zagoriy, V. Mengoli, J. Rojas, K. Jonak et al., 2017 Casein Kinase 1 Coordinates Cohesin Cleavage, Gametogenesis, and Exit from M Phase in Meiosis II. Dev. Cell 40: 37–52.

Attner, M. A., and A. Amon, 2012 Control of the mitotic exit network during meiosis. Mol. Biol. Cell 23: 3122–3132.

Bardin, A. J., and A. Amon, 2001 MEN and SIN: What’s the difference? Nat. Rev. Mol. Cell Biol. 2: 815–826.

Bertazzi, D. T., B. Kurtulmus, and G. Pereira, 2011 The cortical protein Lte1 promotes mitotic exit by inhibiting the spindle position checkpoint kinase Kin4. J. Cell Biol. 193: 1033–1048.

Bizzari, F., and A. L. Marston, 2011 Cdc55 coordinates spindle assembly and chromosome disjunction during meiosis. J. Cell Biol. 193: 1213–1228.

Buonomo, S. B., K. P. Rabitsch, J. Fuchs, S. Gruber, M. Sullivan et al., 2003 Division of the nucleolus and its release of *CDC14* during anaphase of meiosis I depends on separase, *SPO12*, and *SLK19*. Dev Cell 4: 727–739.

Campbell, I. W., X. Zhou, and A. Amon, 2019 The mitotic exit network integrates temporal and spatial signals by distributing regulation across multiple components. eLife 8: e41139.

Chan, L.Y., and A. Amon, 2010 Spindle position is coordinated with cell-cycle progression through establishment of mitotic exit-activating and -inhibitory zones. Mol. Cell 39: 444–454.

Cooper, K. F., M. J. Mallory, D. B. Egeland, M. Jarnik, and R. Strich,2000 Ama1p is a meiosis-specific regulator of the anaphase promoting complex/cyclosome in yeast. Proc. Natl. Acad. Sci. USA 97: 14548–14553.

D’Aquino, K. E., F. Monje-Casas, J. Paulson, V. Reiser, G. M. Charles et al., 2005. The protein kinase Kin4 inhibits exit from mitosis in response to spindle position defects. Mol. Cell 19: 223– 234.

Davidow, L. S., L. Goetsch, and B. Byers, 1980 Preferential occurrence of nonsister spores in two-spored asci of *Saccharomyces cerevisiae*: evidence for regulation of spore-wall formation by the spindle pole body. Genetics 94: 581–595.

Diamond, A. E., J. S. Park, I. Inoue, H. Tachikawa, and A. M. Neiman, 2009 The anaphase promoting complex targeting subunit Ama1 links meiotic exit to cytokinesis during sporulation in *Saccharomyces cerevisiae*. Mol. Biol. Cell 20: 134–145.

Falk, J. E., I. W. Campbell, K. Joyce, J. Whalen, A. Seshan et al., 2016 *LTE1* promotes exit from mitosis by multiple mechanisms. Mol. Biol. Cell 27:3991–4001.

Friesen, H., R. Lunz, S. Doyle, and J. Segall, 1994 Mutation of the *SPS1*-encoded protein kinase of *Saccharomyces cerevisiae* leads to defects in transcription and morphology during spore formation. Genes Dev. 8: 2162–2175.

Gautier, T., T. Berges, D. Tollervey, and E. Hurt, 1997 Nucleolar KKE/D repeat proteins Nop56p and Nop58p interact with Nop1p and are required for ribosome biogenesis. Mol. Cell Biol. 17: 7088–7098.

Gordon, O., C. Taxis, P. J. Keller, A. Benjak, E. H. K. Stelzer et al., 2006 Nud1p, the yeast homolog of Centriolin, regulates spindle pole body inheritance in meiosis. EMBO J. 25: 3856–3868.

Gruneberg, U., K. Campbell, C. Simpson, J. Grindlay, and E. Schiebel, 2000 Nud1p links astral microtubule organization and the control of exit from mitosis. EMBO J. 19: 6475–6488.

Guertin, D. A., L. Chang, F. Irshad, K. L. Gould, and D. McCollum, 2000 The role of the sid1p kinase and cdc14p in regulatin the onset of cytokinesis in fission yeast. EMBO J. 19: 1803–1815.

Hergovich, A., and B. A. Hemmings, 2012 Hippo signaling in the G2/M cell cycle phase: lessons learned from the yeast MEN and SIN pathways. Semin. Cell Dev. Biol. 23: 794–802.

Huang, L. S., H. K. Doherty, and I. Herskowitz, 2005 The Smk1p MAP kinase negatively regulates Gsc2p, a 1,3-beta-glucan synthase, during spore wall morphogenesis in *Saccharomyces cerevisiae*. Proc. Natl. Acad. Sci. USA 102: 12431–12436.

Iwamoto, M. A., S. R. Fairclough, S. A. Rudge, and J. Engebrecht, 2005 Saccharomyces cerevisiae Sps1p regulates trafficking of enzymes required for spore wall synthesis. Eukaryot. Cell 4:536–544.

Jaspersen, S. L., and D. O. Morgan, 2000 Cdc14 activates cdc15 to promote mitotic exit in budding yeast. Curr. Biol. 10: 615–518.

Jones, M. H., J. M. Keck, C. C. Wong, T. Xu, J. R. Yates 3rd et al., 2011 Cell cycle phosphorylation of mitotic exit network (MEN) proteins. Cell Cycle 10: 3435–3440.

Juanes, M. A., and S. Piatti, 2016 The final cut: cell polarity meets cytokinesis at the bud neck in *S. cerevisiae* Cell Mol. Life Sci. 73: 3115–3136.

Kamieniecki, R. J., L. Liu, and D. S. Dawon, 2005 FEAR but not MEN genes are required for exit from meiosis I. Cell Cycle 4: 1093–1098.

Kane, S. M., and R. Roth, 1974 Carbohydrate metabolism during ascospore development in yeast. J. Bacteriol. 118: 8–14.

Kinoshita, E., E. Kinoshita-Kikuta, K. Takiyama, and T. Koike, 2006 Phosphate-binding tag, a new tool to visualize phosphorylated proteins. Mol. Cell. Proteomics 5: 749–757.

Knop, M., and K. Strasser, 2000. Role of the spindle pole body of yeast in mediating assembly of the prospore membrane during meiosis. EMBO J. 19: 3657–3667.

Krishnamoorthy, T., X. Chen, J. Govin, W. L. Cheung, J. Dorsey et al., 2006 Phosphorylation of histone H4 Ser1 regulates sporulation in yeast and is conserved in fly and mouse spermatogenesis. Genes Dev. 20: 2580–2592.

Lam, C., E. Santore, E. Lavoie, L. Needleman, N. Fiacco et al., 2014 A visual screen of protein localization during sporulation identifies new components of prospore membrane-associated complexes in budding yeast. Eukaryot. Cell 13: 383–391.

Lee, B. H., and A. Amon, 2003 Role of Polo-like kinase *CDC5* in programming meiosis I chromosome segregation. Science 300: 482–486.

Lee, S., W. A. Lim, and K. S. Thorn, 2013 Improved blue, green, and red fluorescent protein tagging vectors for S. cerevisiae. PLoS One 8: e67902.

Longtine, M. S., A. McKenzie, D. J. Demarini, N. G. Shah, A. Wach et al., 1998 Additional modules for versatile and economical PCR-based gene deletion and modification in *Saccharomyces cerevisiae*. Yeast 14: 953–961.

Luca, F. C., M. Mody, C. Kurischko, D. M. Roof, T. H. Giddings, et al., 2001 *Saccharomyces cerevisiae* Mob1p is required for cytokinesis and mitotic exit. Mol. Cell Biol. 21: 6972–6983.

Maekawa, H., C. Priest, J. Lechner, G. Pereira, and E. Schiebel, 2007 The yeast centrosome translates the positional information of the anaphase spindle into a cell cycle signal. J. Cell Biol. 179: 423–436.

Mah, A. S., J. Jang, and R. J. Deshaies, 2001 Protein kinase Cdc15 activates the Dbf2-Mob1 kinase complex. Proc. Natl. Acad. Sci. USA 98: 7325–7330.

Maier, P., N. Rathfelder, M.G. Finkbeiner, C. Taxis, M. Mazza et al., 2007 Cytokinesis in yeast meiosis depends on the regulated removal of Ssp1p from the prospore membrane. EMBO J. 26: 1843–1852.

Manzoni, R., F. Montani, C. Visintin, F. Caudron, A. Ciliberto et al., 2010 Oscillations in Cdc14 release and sequestration reveal a circuit underlying mitotic exit. J. Cell Biol. 190: 209–222.

Marston, A. L., B. H. Lee, and A. Amon, 2003 The Cdc14 phosphatase and the FEAR network control meiotic spindle disassembly and chromosome segregation. Dev. Cell 4: 711–726.

Meitinger, F., M. E. Boehm, A. Hofmann, B, Hub, H. Zentgraf et al., 2011 Phosphorylation-dependent regulation of the F-BAR protein Hof1 during cytokinesis. Genes Dev. 25: 875–888.

Mohl, D. A., M. J. Huddleston, T. S. Collingwood, R. S. Annan, and R. J. Deshaies, 2009 Dbf2-Mob1 drives relocalization of protein phosphatase Cdc14 to the cytoplasm during exit from mitosis. J. Cell Biol. 184: 527–539.

Moreno-Borchart, A. C., K. Strasser, M. G. Finkbeiner, A. Shevchenko, A. Shevchenko et al., 2001 Prospore membrane formation linked to the leading edge protein (LEP) coat assembly. EMBO J. 20: 6946–6957.

Nakanishi, H., P. de los Santos, and A. M. Neiman, 2004 Positive and negative regulation of a SNARE protein by control of intracellular localization. Mol. Biol. Cell 15: 1802–1815.

Neiman, A.M., L. Katz, and P. J Brennwald, 2000 Identification of Domains Required for Developmentally Regulated SNARE Function in *Saccharomyces cerevisiae*. Genetics 155: 1643–1655.

Neiman, A. M., 2011 Sporulation in the budding yeast *Saccharomyces cerevisiae*. Genetics 189: 737–765.

Nickas, M. E., A. E. Diamond, M.-J. Yang, and A. M. Neiman, 2004 Regulation of spindle pole function by an intermediary metabolite. Mol. Biol. Cell 15: 2606–2616.

Oh, Y., K. J. Chang, P. Orlean, C. Wloka, R. Deshaies et al., 2012 Mitotic exit kinase Dbf2 directly phosphorylates chitin synthase Chs2 to regulate cytokinesis in budding yeast. Mol. Biol. Cell 23: 2445–2456.

Pablo-Hernando, M. E., Y. Arnaiz-Pita, H. Nakanishi, D. Dawson, F. del Rey, et al., 2007 Cdcc15 is required for spore morphogenesis independently of Cdc14 in *Saccharomyces cerevisiae*. Genetics 177: 281–293.

Paulissen, S.M., C.J. Slubowski, J.M. Roesner, and L.S. Huang, 2016 Timely Closure of the Prospore Membrane Requires *SPS1* and *SPO77* in *Saccharomyces cerevisiae*. Genetics 204: 1203–1216.

Pereira, G., and E Schiebel, 2005 Kin4 kinase delays mitotic exit in response to spindle alignment defects. Mol. Cell 19: 209–221.

Philips, J., and I. Herskowitz, 1998 Identification of Kel1p, a Kelch domain-containing protein involved in cell fusion and morphology in *Saccharomyces cerevisiae*. J. Cell Biol. 143: 375–398.

Poon, C. L. C., W. Liu, Y. Song, M. Gomez, Y. Kulaberoglu et. al., 2018 A Hippo-like Signaling Pathway Controls Tracheal Morphogenesis in *Drosophila melanogaster*. Dev Cell 47: 564–575.

Rabitsch, K.P., A. Toth, M. Galova, A. Schleiffer, G. Schaffner et al., 2001 A screen for genes required for meiosis and spore formation based on whole-genome expression. Curr Biol. 11: 1001–1009.

Renicke, C., A.-K. Allman, A. P. Lutz, T. Heimerl, and C. Taxis, 2017 The Mitotic Exit Network regulates spindle pole body selection during sporulation of *Saccharomyces cerevisiae*. Genetics 206: 919–937.

Rock, J. M., and A. Amon, 2011 Cdc15 integrates Tem1 GTPase-mediated spatial signals with Polo kinase-mediated temporal cues to activate mitotic exit. Genes Dev. 25: 1943–1954.

Rock, J. M., D. Lim, L. Stach, R. W. Ogrodowicz, J. M. Keck et al., 2013 Activation of the yeast Hippo pathway by phosphorylation-dependent assembly of signaling complexes. Science 340: 871–875.

Rose, M. D., and G. R. Fink, 1990 Methods in Yeast Genetics, Cold Spring Harbor, NY: Cold Spring Harbor Laboratory Press.

Schindelin, J., I. Arganda-Carreras, E. Frise, V. Kaynig, M. Longair et al., 2012 Fiji: an open-source platform for biological-image analysis. Nat Methods 9: 676–82.

Schneider, C. A., W. S. Rasband, and K.W. Eliceiri, 2012 NIH Image to ImageJ: 25 years of image analysis. Nat Methods 9: 671–675.

Schweitzer, B., and P. Philippsen, 1991 *CDC15*, an essential cell cycle gene in *Saccharomyces cerevisiae*, encodes a protein kinase domain. Yeast 7: 265–273.

Shou, W., J. H. Seol, A. Shevchenko, C. Baskerville, D. Moazed et al., 1999 Exit from mitosis is triggered by Tem1-dependent relase of the protein phosphatase Cdc14 from nucleolar RENT complex. Cell 97: 233–244.

Slubowski, C.J., A.D. Funk, J.M. Roesner, S.M. Paulissen, and L.S. Huang, 2015 Plasmids for C-terminal tagging in *Saccharomyces cerevisiae* that contain improved GFP proteins, Envy and Ivy. Yeast 32: 379–387.

Slubowski, C.J., S.M. Paulissen, and L.S. Huang, 2014 The GCKIII kinase Sps1 and the 14-3-3 isoforms, Bmh1 and Bmh2, cooperate to ensure proper sporulation in *Saccharomyces cerevisiae*. PLoS One 9: e113528.

Stegmeier, F., and A. Amon, 2004 Closing mitosis: the functions of the Cdc14 phosphatase and its regulation. Annu. Rev. Genet. 38: 203–232.

Taxis, C., P. Keller, Z. Kavagiou, L. J. Jensen, J. Colombelli et al., 2005 Spore number control and breeding in *Saccharomyces cerevisiae*: a key role for a self-organizing system. J. Cell Biol. 171: 627–640.

Visintin, R., K. Craig, E.S. Hwang, S. Prinz, M. Tyers et al., 1998 The phosphatase Cdc14 triggers mitotic exit by reversal of Cdk-dependent phosphorylation. Mol. Cell 2: 709–718.

Visintin R., and A. Amon, 2001 Regulation of the mitotic exit protein kinases Cdc15 and Dbf2. Mol. Biol. Cell 12: 2961–2974.

Weiss, E. L., 2012 Mitotic exit and separation of mother and daughter cells. Genetics 192: 1165–1202.

Whinston, E., G. Omerza, A. Singh, C. W. Tio, and E. Winter, 2013 Activation of the Smk1 mitogen-activated protein kinase by developmentally regulated autophosphorylation. Mol. Cell. Biol. 33: 688–700.

