## Supplemental Figures and Tables for "A non-canonical Hippo pathway regulates spindle disassembly and cytokinesis during meiosis in *Saccharomyces cerevisiae*"

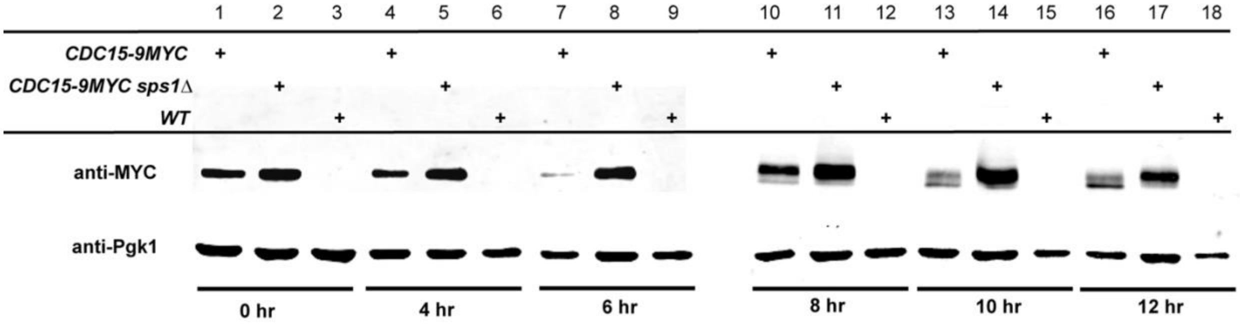


**Figure S1.** *SPS1* is not required for Cdc15 phosphorylation. Wild type (WT: LH177) and strains containing *CDC15-9myc* (LH1070) and *CDC15-9myc* in the *sps1Δ* strain (LH1083) were sporulated and lysates were prepared for immunoblotting. Lanes 1-9 were run on the same SDS-PAGE gel; Lanes 10-18 are on the same SDS-PAGE gel. Pgk1 was used as a loading control.

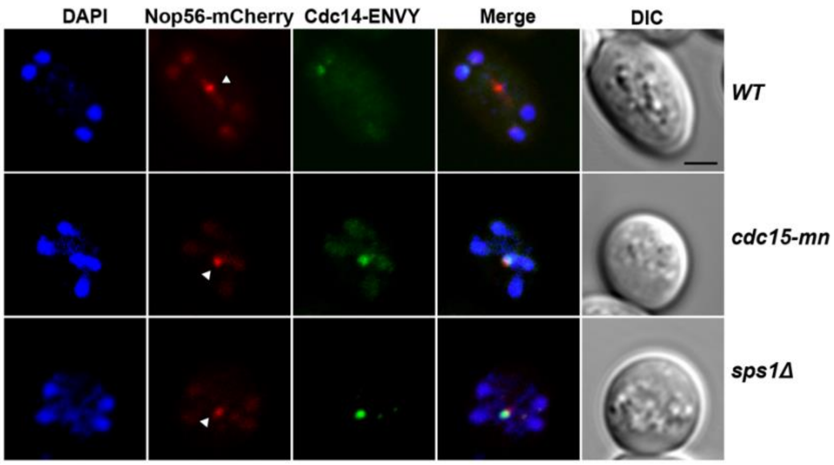


**Figure S2.** Cdc14 remains in the nucleolus in *cdc15* and *sps1* mutants. In anaphase II, Cdc14-ENVY co-localizes with the nucleolar marker Nop56 in *cdc15-mn* (LH1086) and *sps1Δ* (LH1085), but is released from the nucleolus in wild type cells (LH1084). White arrowhead points to nucleolus. Scale bar = 2 μm.

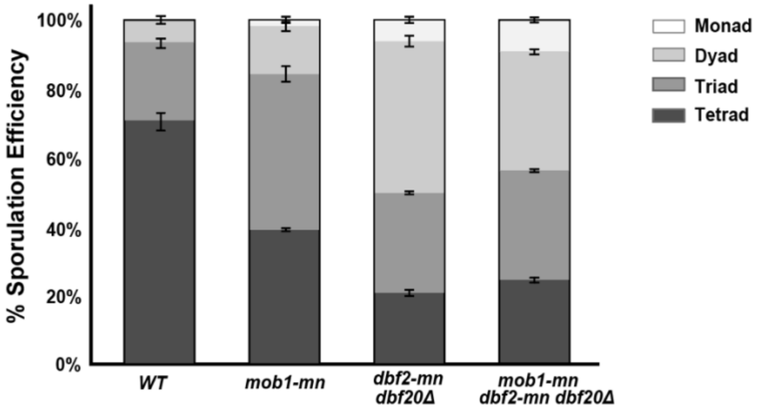


**Figure S3.** *MOB1*, *DBF2*, and *DBF20* affect spore number control. The number of spores packaged was assayed by counting refractile spores in wild type (LH902), *mob1-mn* (LH1087), *dbf2-mn dbf20Δ* (LH1088), *mob1-mn, dbf2-mn, dbf20Δ* (LH1068). Monads have 1 spore per ascus, dyads have 2, triads have 3 and tetrads have 4. Error bars depict the standard error of the mean. All three mutant strains are significantly different from wild type, but not from one another, in the fraction of mutant spores formed (one-way ANOVA [$F(3,8)=64$, $p<0.001$], followed by Tukey HSD post hoc test (alpha = 0.01)).

**TABLE S1: Yeast Strains**

| Strain | Genotype | Source |
|---|---|---|
| A20239 | MATa/MATα ho::LYS2/ ho::LYS2 lys2/ lys2 leu2::hisG/ leu2::hisG his3::hisG/ his3::hisG trp1::hisG/ trp1::hisG ura3::pGPD1GAL4(848).ER:URA3/ ura3::pGPD1-GAL4(848).ER:URA3 GALNDT80:TRP1/ GAL-NDT80:TRP KanMX6:pCLB2-3HA-MOB1/KanMX6:pCLB2-3HA-MOB1 | Attner et al. 2012 |
| A22416 | MATa/MATα ho::LYS2/ ho::LYS2 lys2/ lys2 leu2::hisG/ leu2::hisG his3::hisG/ his3::hisG trp1::hisG/ trp1::hisG ura3::pGPD1GAL4(848).ER:URA3/ ura3::pGPD1-GAL4(848).ER:URA3 GALNDT80:TRP1/ GAL-NDT80:TRP1 CDC15-9MYC:TRP1/CDC15-9MYC:TRP1 | Attner et al. 2012 |
| HI50 | MATa/MATα ura3/ura3 leu2/leu2 trp1::hisG/trp1::hisG ARG4/arg4-NspI his3ΔSK/ his3ΔSK hoΔ::LYS2/hoΔ::LYS2 lys2/lys2 RME1/rme1Δ:: LEU2 mxKAN:prCLB2:HA:CDC15/mxKAN:prCLB2:HA:CDC15 | Pablo-Hernando et al. 2007 |
| AN117-4B | MATα his3 ura3 trp1::hisG leu2 arg4-NspI lys2 hoΔ::LYS2 rme1::LEU2 | Neiman et al. 2000 |
| YS429 | MATa/MATα hoΔ::LYS2/hoΔ::LYS2 lys2/lys2 ura3/ura3 leu2/leu2 trp1::hisG/trp1::hisG his3/his3 arg4-Nsp1/ARG4 RME1/rme1Δ::LEU2 mxKAN:prCLB2:HA:DBF2/mxKAN:prCLB2:HA:DBF2 dbf20Δ::kanMX6/ dbf20Δ::kanMX6 | This study |
| LH177 | MATa/MATα ho::LYS2/ho::LYS2 lys2/lys2 ura3/ura3 leu2/leu2 his3/his3 trp1ΔFA/trp1ΔFA | Huang et al. 2005 |
| LH872 | MATa/MATα ho::LYS2/ho::LYS2 lys2/lys2 ura3/ura3 leu2/leu2 his3/his3 trp1ΔFA/trp1ΔFA sps1::LEU2$_{C.g.}$/sps1::LEU2$_{C.g.}$ | Slubowski et al. 2014 |
| LH875 | MATa/MATα ho::hisG/ho::hisG lys2/lys2 ura3/ura3 leu2/leu2 his3/his3 trp1ΔFA/trp1ΔFA SPS1-13x myc-TRP1/SPS1-13x myc-TRP1 | Slubowski et al. 2014 |
| LH902 | MATa/MATα ho::LYS2/ho::LYS2 lys2/lys2 ura3/ura3 leu2/leu2 his3/his3 trp1ΔFA/trp1ΔFA HTB2:mCherry:TRP1$_{C.g.}$/HTB2:mCherry:TRP1$_{C.g.}$ | Parodi et al. 2012 |
| LH966 | MATa/MATα ho::hisG/ho::hisG lys2/lys2 ura3/ura3 leu2/leu2 his3/his3 trp1ΔFA/trp1ΔFA HTB2-mCherry-TRP1$_{C.g.}$/HTB2-mCherry-TRP1$_{C.g.}$ sps1::LEU2$_{C.g.}$/sps1::LEU2$_{C.g.}$ | Slubowski et al. 2014 |
| LH976 | MATa/MATα ho::LYS2/ho::LYS2 lys2/lys2 ura3/ura3 leu2/leu2 his3/his3 trp1ΔFA/trp1ΔFA sps1::HIS3$_{C.g.}$/sps1::HIS3$_{C.g.}$ HTB2:mCherry:TRP1$_{C.g.}$/HTB2:mCherry:TRP1$_{C.g.}$ | Slubowski et al. 2014 |
| LH986 | MATa/MATα ho::LYS2/ho::LYS2 lys2/lys2 ura3/ura3 leu2/leu2 his3/his3 trp1ΔFA/trp1ΔFA sfGFP:SPS1/sfGFP:SPS1 HTB2:mCherry:TRP1$_{C.g.}$/HTB2:mCherry:TRP1$_{C.g.}$ | Slubowski et al. 2014 |
| LH1010 | MATa/MATα ho::LYS2/ho::LYS2 lys2/lys2 ura3/ura3 leu2/leu2 his3/his3 trp1ΔFA/trp1ΔFA spo77::HIS3$_{C.g.}$/ spo77::HIS3$_{C.g.}$ HTB2:mCherry:TRP1$_{C.g.}$/HTB2:mCherry:TRP1$_{C.g.}$ | Paulissen et al. 2016 |
| LH1014 | MATa/MATα ho::LYS2/ho::LYS2 lys2/lys2 ura3/ura3 leu2/leu2 his3/his3 trp1ΔFA/trp1ΔFA ama1::TRP1$_{C.g.}$/ama1::TRP1$_{C.g.}$ HTB2:mCherry:TRP1$_{C.g.}$/HTB2:mCherry:TRP1$_{C.g.}$ | Paulissen et al. 2016 |
| LH1066 | MATa/MATα ho::LYS2/ho::LYS2 lys2/lys2 ura3/ura3 leu2/leu2 his3/his3 trp1ΔFA/trp1ΔFA mxKAN:prCLB2:HA:CDC15/mxKAN:prCLB2:HA:CDC15 HTB2:mCherry:URA3$_{K.l.}$/HTB2:mCherry:URA3$_{K.l.}$ | This study |
| LH1067 | MATa/MATα ho::LYS2/ho::LYS2 lys2/lys2 ura3/ura3 leu2/leu2 his3/his3 trp1ΔFA/trp1ΔFA mxKAN:prCLB2:HA:CDC15/mxKAN:prCLB2:HA:CDC15 sps1::LEU2$_{C.g.}$/sps1::LEU2$_{C.g.}$ HTB2:mCherry:URA3$_{K.l.}$/HTB2:mCherry:URA3$_{K.l.}$ | This study |

**TABLE S1, continued**

| Strain | Genotype | Source |
|---|---|---|
| LH1068 | *MAT**a**/MATα ho::LYS2/ho::LYS2 lys2/lys2 ura3/ura3 leu2/leu2 his3/his3 trp1ΔFA/trp1ΔFA KanMX6:pCLB2-3HA-MOB1/KanMX6:pCLB2-3HA-MOB1 mxKAN:prCLB2:HA:DBF2/mxKAN:prCLB2:HA:DBF2 dbf20Δ::kanMX6/dbf20Δ::kanMX6 HTB2-mCherry-TRP1*$_{C.g.}$*/HTB2-mCherry-TRP1*$_{C.g.}$ | This study |
| LH1069 | *MAT**a**/MATα ho::LYS2/ho::LYS2 lys2/lys2 ura3/ura3 leu2/leu2 his3/his3 trp1ΔFA/trp1ΔFA mxKAN:prCLB2:HA:CDC15/mxKAN:prCLB2:HA:CDC15 SPS1:13myc:TRP1/SPS1:13myc:TRP1* | This study |
| LH1070 | *MAT**a**/MATα ho::LYS2/ho::LYS2 lys2/lys2 ura3/ura3 leu2/leu2 his3/his3 trp1ΔFA/trp1ΔFA CDC15:9myc:TRP1/CDC15:9myc:TRP1 HTB2:mCherry:TRP1*$_{C.g.}$*/HTB2:mCherry:TRP1*$_{C.g.}$ | This study |
| LH1071 | *MAT**a**/MATα ho::LYS2/ho::LYS2 lys2/lys2 ura3/ura3 leu2/leu2 his3/his3 trp1ΔFA/trp1ΔFA sfGFP:SPS1/sfGFP:SPS1 CDC15:9myc:TRP1/CDC15:9myc:TRP1 HTB2:mCherry:TRP1*$_{C.g.}$*/HTB2:mCherry:TRP1*$_{C.g.}$ | This study |
| LH1072 | *MAT**a**/MATα ho::LYS2/ho::LYS2 lys2/lys2 ura3/ura3 leu2/leu2 his3/his3 trp1ΔFA/trp1ΔFA mxKAN:prCLB2:HA:CDC15/mxKAN:prCLB2:HA:CDC15 HTB2:mCherry:URA3*$_{K.l.}$*/HTB2:mCherry:URA3*$_{K.l.}$ | This study |
| LH1075 | *MAT**a**/MATα ho::LYS2/ho::LYS2 lys2/lys2 ura3/ura3 leu2/leu2 his3/his3 trp1ΔFA/trp1ΔFA mxKAN:prCLB2:HA:CDC15/mxKAN:prCLB2:HA:CDC15 ama1::TRP1*$_{C.g.}$*/ama1::TRP1*$_{C.g.}$ | This study |
| LH1077 | *MAT**a**/MATα ho::LYS2/ho::LYS2 lys2/lys2 ura3/ura3 leu2/leu2 his3/his3 trp1ΔFA/trp1ΔFA HTB2:mCherry:TRP1*$_{C.g.}$*/HTB2:mCherry:TRP1*$_{C.g.}$ *CDC14:ENVY:HIS3*$_{S.p.}$*/ +* | This study |
| LH1078 | *MAT**a**/MATα ho::LYS2/ho::LYS2 lys2/lys2 ura3/ura3 leu2/leu2 his3/his3 trp1ΔFA/trp1ΔFA HTB2:mCherry:TRP1*$_{C.g.}$*/HTB2:mCherry:TRP1*$_{C.g.}$ *CDC14:ENVY:HIS3*$_{S.p.}$*/ + sps1::LEU2* $_{C.g.}$*/sps1::LEU2* $_{C.g.}$ | This study |
| LH1079 | *MAT**a**/MATα ho::LYS2/ho::LYS2 lys2/lys2 ura3/ura3 leu2/leu2 his3/his3 trp1ΔFA/trp1ΔFA HTB2:mCherry:TRP1*$_{C.g.}$*/HTB2:mCherry:TRP1*$_{C.g.}$ *CDC14:ENVY:HIS3*$_{S.p.}$*/ + mxKAN:prCLB2:HA:CDC15/mxKAN:prCLB2:HA:CDC15* | This study |
| LH1080 | *MAT**a**/MATα ho::LYS2/ho::LYS2 lys2/lys2 ura3/ura3 leu2/leu2 his3/his3 trp1ΔFA/trp1ΔFA HTB2:mCherry:TRP1*$_{C.g.}$*/HTB2:mCherry:TRP1*$_{C.g.}$ *CDC14:ENVY:HIS3*$_{S.p.}$*/ + KanMX6:pCLB2-3HA-MOB1/KanMX6:pCLB2-3HA-MOB mxKAN:prCLB2:HA:DBF2/ mxKAN:prCLB2:HA:DBF2 dbf20Δ::kanMX6/dbf20Δ::kanMX6* | This study |
| LH1081 | *MAT**a**/MATα ho::LYS2/ho::LYS2 lys2/lys2 ura3/ura3 leu2/leu2 his3/his3 trp1ΔFA/trp1ΔFA KanMX6:pCLB2-3HA-MOB1/KanMX6:pCLB2-3HA-MOB1 mxKAN:prCLB2:HA:DBF2/ mxKAN:prCLB2:HA:DBF2 dbf20Δ::kanMX6/dbf20Δ::kanMX6* | This study |
| LH1083 | *MAT**a**/MATα ho::LYS2/ho::LYS2 lys2/lys2 ura3/ura3 leu2/leu2 his3/his3 trp1ΔFA/trp1ΔFA CDC15:9myc:TRP1/CDC15:9myc:TRP1 HTB2:mCherry:TRP1*$_{C.g.}$*/HTB2:mCherry:TRP1*$_{C.g.}$ *sps1::LEU2* $_{C.g.}$*/ sps1::LEU2* $_{C.g.}$ | This study |
| LH1084 | *MAT**a**/MATα ho::LYS2/ho::LYS2 lys2/lys2 ura3/ura3 leu2/leu2 his3/his3 trp1ΔFA/trp1ΔFA CDC14:ENVY: HIS3*$_{S.p.}$*/ CDC14:ENVY: HIS3*$_{S.p}$ *NOP56:mCherry: TRP1*$_{C.g.}$*/ NOP56:mCherry: TRP1* $_{C.g.}$ | This study |

**TABLE S1, continued**

| Strain | Genotype | Source |
|---|---|---|
| LH1085 | *MAT**a**/MATα ho::LYS2/ho::LYS2 lys2/lys2 ura3/ura3 leu2/leu2 his3/his3 trp1ΔFA/trp1ΔFA CDC14:ENVY:HIS3$_{S.p.}$/ CDC14:ENVY:HIS3$_{S.p.}$ sps1::LEU2 $_{C.g.}$/sps1::LEU2 $_{C.g.}$ NOP56:mCherry: TRP1$_{C.g.}$/ NOP56:mCherry: TRP1 $_{C.g.}$* | This study |
| LH1086 | *MAT**a**/MATα ho::LYS2/ho::LYS2 lys2/lys2 ura3/ura3 leu2/leu2 his3/his3 trp1ΔFA/trp1ΔFA CDC14:ENVY:HIS3$_{S.p.}$/CDC14:ENVY:HIS3$_{S.p.}$ mxKAN:prCLB2:HA:CDC15/mxKAN:prCLB2:HA:CDC15 NOP56:mCherry: TRP1$_{C.g.}$/ NOP56:mCherry: TRP1 $_{C.g.}$* | This study |
| LH1087 | *MAT**a**/MATα ho::LYS2/ho::LYS2 lys2/lys2 ura3/ura3 leu2/leu2 his3/his3 trp1ΔFA/trp1ΔFA KanMX6:pCLB2-3HA-MOB1/KanMX6:pCLB2-3HA-MOB HTB2:mCherry:TRP1$_{C.g.}$/HTB2:mCherry:TRP1$_{C.g.}$* | This study |
| LH1088 | *MAT**a**/MATα ho::LYS2/ho::LYS2 lys2/lys2 ura3/ura3 leu2/leu2 his3/his3 trp1ΔFA/trp1ΔFA mxKAN:prCLB2:HA:DBF2/mxKAN:prCLB2:HA:DBF2 dbf20Δ::kanMX6/dbf20Δ::kanMX6 HTB2:mCherry:TRP1$_{C.g.}$/ HTB2:mCherry:TRP1$_{C.g.}$* | This study |

**TABLE S2: Yeast Strains Containing Plasmids**

| Strain | Genotype | Source |
|---|---|---|
| LH917 | LH902 plus pRS426-G20 | Parodi *et al*. 2012 |
| LH1047 | LH976 (*sps1Δ*) plus pRS426-G20 | Paulissen *et al*. 2016 |
| LH1073 | LH1072 (*cdc15-mn*) plus pRS426-E20 | This study |
| LH1074 | LH1067 (*sps1 Δ cdc15-mn*) plus pRS426-E20 | This study |
| LH1076 | LH1075 (*cdc15-mn ama1 Δ*) plus pRS426-E20 | This study |
| LH1081 | LH902 plus pRS426-E20 | This study |
| LH1082 | LH1068 (*mob1-mn dbf2-mn dbf20 Δ*) plus pRS426-E20 | This study |
| LH1089 | LH976 (*sps1 Δ*) plus pRS426-E20 | This study |

**TABLE S3: Genes screened for H4S1p phenotype**

| Gene Name | | | | |
|---|---|---|---|---|
| *ADY2* | *HIT1* | *PLP1* | *TMA64* | *YDL180W* |
| *ADY3* | *HST4* | *PRD1* | *TOS10* | *YDL186W* |
| *ALK2* | *HUL5* | *PRM2* | *TOS8* | *YDR015C* |
| *AMA1* | *ISC10* | *PRR2* | *TPH3* | *YDR018C* |
| *AMA1* | *LAA2* | *RGD1* | *TRS85/GSG1*[2] | *YDR042C* |
| *AOR1* | *LCD1* | *RRG1* | *UBP1* | *YDR250C* |
| *API2* | *MAM1* | *RRT5* | *ULI1* | *YDR491C* |
| *ATG9*[2] | *MCH1* | *RTT12* | *VHS1* | *YEL023C* |
| *ATG18*[2] | *MCY1* | *RXT3* | *YAP6* | *YER076C* |
| *AZR1* | *MFG1* | *SET1*[2] | *YCK3* | *YER084W* |
| *CDC15*[1, 2] | *MGR1* | *SPO22* | *YEF1* | *YER085C* |
| *CSM1* | *MIP6* | *SPO23* | *YSP2* | *YFL040W* |
| *DCV1* | *MND2*[2] | *SPO71* | *YSW1* | *YFR018C* |
| *DIG2* | *MPS3* | *SPO73* | *YUH1* | *YGL081W* |
| *DMA1* | *MUS81* | *SPO75* | *ZRG8* | *YGL138C* |
| *DTR1* | *NKP1* | *SPO77*[2] | *YBR063C* | *YGL230C* |
| *EDC2* | *NPP2* | *SPR6* | *YBR134W* | *YGL242C* |
| *FMP10* | *OSW7* | *SPR28* | *YBR144C* | *YGR016W* |
| *FNH1* | *PES4* | *SPS22* | *YBR184W* | *YIL024C* |
| *GAT4* | *PET18* | *SSM4* | *YBR232C* | *YIL055C* |
| *GIP1* | *PEX22* | *SSP2* | *YCR061W* | *YJR037W* |
| *GIS1* | *PEX28* | *SWM1* | *YDL109C* | |
| *GMC1* | *PHO92* | *SYP1* | *YDL114W* | |
| *HIM1* | *PKP1* | *THI74* | *YDL162C* | |

[1]All strains screened were from the yeast gene knock-out collection in the SK1 background from Rabitsch *et al*. 2001 except for the *cdc15* mutant containing strain, which was from HI50 (Pablo-Hernando *et al*. 2007).

[2]The *atg9, atg18*, *cdc15*, *mnd2*, *set1, spo77* and *trs85* containing mutant strains exhibited decreased H4S1p phosphorylation.
